# Sub-Optimal Learning of Tactile-Spatial Predictions in Patients with Complex Regional Pain Syndrome

**DOI:** 10.1101/775676

**Authors:** Christopher A. Brown, Ingrid Scholtes, Nicholas Shenker, Michael C. Lee

**Affiliations:** Department of Psychological Sciences, University of Liverpool, Bedford Street South, Liverpool L69 7ZA, UK; University Division of Anaesthesia, University of Cambridge, Addenbrooke’s Hospital, Hills Road, Cambridge CB2 0QQ, UK; Department of Rheumatology, Addenbrooke’s Hospital, Hills Road, Cambridge CB2 0QQ, UK

## Abstract

In Complex Regional Pain Syndrome (CRPS), tactile sensory deficits have motivated the therapeutic use of sensory discrimination training. However, the hierarchical organisation of the brain is such that low-level sensory processing can be dynamically influenced by higher-level knowledge, e.g. knowledge learnt from statistical regularities in the environment. It is unknown whether the learning of such statistical regularities is impaired in CRPS. Here, we employed a hierarchical Bayesian model of predictive coding to investigate statistical learning of tactile-spatial predictions in CRPS. Using a sensory change-detection task, we manipulated bottom-up (spatial displacement of a tactile stimulus) and top-down (probabilistic structure of occurrence) factors to estimate hierarchies of prediction and prediction error signals, as well as their respective precisions or reliability. Behavioural responses to spatial changes were influenced by both the magnitude of spatial displacement (bottom-up) and learnt probabilities of change (top-down). The Bayesian model revealed that patients’ predictions (of spatial displacements) were found to be less precise, deviating further from the ideal (statistical optimality) compared to healthy controls. This imprecision was less context-dependent, i.e. more enduring across changes in probabilistic context and less finely-tuned to statistics of the environment. This caused greater precision on prediction errors, resulting in predictions that were driven more by momentary spatial changes and less by the history of spatial changes. These results suggest inefficiencies in higher-order statistical learning in CRPS. This may have implications for therapies based on sensory re-training whose effects may be more short-lived if success depends on higher-order learning.

## Introduction

Complex regional pain syndrome (CRPS) is characterised by disproportionate pain that is usually initiated by peripheral trauma to a limb (Birklein et al., 2018). The later stages of the disorder are characterised mainly by intractable pain that is poorly understood. Numerous behavioural studies have revealed links between pain and deficits in spatial discrimination of tactile stimuli (Förderreuther et al., 2004; Lewis and Schweinhardt, 2012; Pleger et al., 2006). Those studies, coupled with initial findings of altered somatotopic mapping of the affected (painful) limb in the primary somatosensory cortex (S1) in CRPS (Di Pietro et al., 2013; Kuttikat et al., 2016a), have motivated therapies focussed on S1 re-mapping, for example tactile discrimination training (Moseley et al., 2008). However, more recent high-resolution functional Magnetic Resonance Imaging (fMRI) studies have failed to replicate somatotopic abnormalities of S1 (Di Pietro et al., 2015; Mancini et al., 2018; van Velzen et al., 2016). This has led to further hypotheses of deficits in higher-order mechanisms to explain sensory symptoms in CRPS (Kuttikat et al., 2016a; Popkirov et al., 2018).

Perceptual decision-making is shaped by both afferent feedforward (somatosensory) and cortical feedback mechanisms (Allen et al., 2015; Auksztulewicz et al., 2012; Hegner et al., 2017; Langner et al., 2011). The hierarchical organisation of the brain is such that low-level sensory processing can be dynamically influenced, via feedback mechanisms, by higher-level knowledge (de Lange et al., 2018). In particular, statistical regularities in the environment, i.e. likelihood of event occurrence, can be learnt over time to aid sensory detection or discrimination (Hasson, 2017). In the human brain, cortical anatomy is hierarchically organised such as to reflect the multiple temporal scales at which environmental states can evolve (Kiebel et al., 2008). Hierarchical predictive coding (HPC) is a model of how feedforward and feedback loops contribute to perceptual inference by utilising information at higher spatial and temporal scales (Bastos et al., 2012; Bogacz, 2017; Friston, 2018; Friston and Kiebel, 2009; Rao and Ballard, 1999). HPC explains how the brain might reduce redundancy (and increase efficiency) by only propagating unpredicted (e.g. novel or surprising) sensory inputs or information from low-level to high-level cortical regions. Unpredicted information is propagated forwards as prediction errors (PEs), namely the mismatch or difference between predicted and actual input at each level in the cortical hierarchy.

From the perspective of HPC, in order to process sensory information efficiently, the brain aims to minimise prediction error (Friston & Kiebel, 2009). Persistent prediction errors might be caused by impaired learning from sensory information (Bogacz, 2017), including from the statistics of the environment, i.e. the probability of a change in sensory input and those probabilities evolve over time (environmental volatility). Furthermore, computational modelling has revealed that behavioural and physiological indices of sensory perception in healthy individuals conform to Bayesian models of hierarchical predictive coding, including the Hierarchical Gaussian Filter (HGF) (Mathys et al., 2014). Such Bayesian models consider predictions and PEs as probability distributions characterised by means (magnitude) and precisions (inverse variance, or reliability). The balance of precision between PEs and predictions determines the extent to which bottom-up information (represented by PE) is used to update predictions; i.e. it is an indicator of need of further learning (Bogacz, 2017).

Using the HGF, we sought to test the novel hypothesis that patients with CRPS are sub-optimal in their learning from tactile-spatial events that normally enable efficient prediction error minimisation. By applying this model to response times from a sensory change-detection paradigm, here we show reduced efficiency in predictive coding of spatial information in patients with CRPS.

## MATERIALS AND METHODS

### Study design and setting

The study was observational in nature and employed a case-control design. Two groups of participants were recruited (CRPS and Healthy Controls, HC). All participants attended an experimental session at the Clinical Research Facility at Cambridge University Hospitals NHS Trust (CUH) to complete all study tasks reported below. Ethical approval for the research was obtained by the London (Bromley) ethics committee (reference number 15-LO-1624). The recruitment period was between September 2016 and April 2018.

### Participants

Details of recruitment, inclusion/exclusion criteria and characteristics of the participants (including controls) are in Supplementary Materials. In summary, 22 patients were recruited with a mixture of upper and lower limb CRPS (Supplementary Tables 1 and 2). There were 22 healthy controls (HC) included in the analysis. There was an average age difference between groups with medians of 57 (CRPS) and 40.5 (HC). This potential confound was addressed by selection of n=15 per group for all group effect analyses, which was the maximum number that allowed a close age-matching between groups: For the CRPS group, who were older on average, the 7 oldest patients were excluded so that the remaining oldest patient was age-matched to the oldest HC participant; for HC group, a participant was selected if their age matched most closely with a patient in the CRPS group, such that each CRPS patient was paired with the closest possible HC participant. This resulted in median ages of 50 (CRPS) and 49 (HC), with similar numbers of females in each group (80% and 73.3% respectively). In the CRPS subgroup, the median symptom duration was 7 years. All participants were right-handed, did not have any current or previous diagnosis of peripheral neuropathy, stroke, transient ischemic attack, multiple sclerosis, malignancy or seizure. The participants were required to refrain from consuming alcohol or smoking tobacco for 24 hours and caffeine for 12 hours prior to the study. All participants signed an informed consent form prior to taking part.

### Tactile Spatial Oddball Task (TSOT)

The TSOT was programmed in Matlab. The outputs of the programme were to a Labjack U3-HV which sent digital outputs to a Digitimer DS7A Electrical Stimulator. The DS7A was set with a pulse width of 200µs. The DS7A sent electrical outputs to a Digitimer D188 Electrode Selector, which contains an electronic switch to select outputs to one of eight pairs of stimulating digital ring electrodes. Each pair of stimulating electrodes consisted of an anode placed on the near-side of the knuckle (closest to the wrist), and the cathode (black wire) on the far-side (closest to the finger-tips), spaced apart by 3cm, which was consistent between electrode pairs on each digit. The PC was also connected to a foot pedal via USB for participant responses.

The intensity of electrical stimulation for each participant was based on sensory detection threshold testing using an adaptive staircase. The initial stimulus was 1mA, with increment of 0.2 mA for each consecutive trial until the participant reports a sensation, after which the current was decreased by 0.1mA for subsequent trials until loss of sensation occurs. The current used in the preceding trial at which the participant did feel a sensation was noted. The participant was asked to report the first time they felt any sensation on the ring finger of the hand of the unaffected side; this procedure was then repeated for the affected side. Each stimulus occurred at a random and unexpected time (between 1 and 3s after responding to the previous trial) to minimise effects of temporal anticipation.

We employed electrical stimuli at 3x sensory detection threshold for the experiment to ensure that each stimulus was clearly perceived. The stimuli were entirely painless for healthy controls. For those patients who found the stimulus uncomfortable at that level (8 out of 22 patients), the current was reduced by 0.1mA at a time until the stimulus was just below the threshold for pain. There were no group differences in the currents used during the task; we also explored the relationship between current delivered and key dependent variables from the study (see Supplementary Materials for results).

Participants were seated in a comfortable chair. Digital ring electrode pairs were placed on each digit of all four fingers of each hand (excluding the thumb). Tissue paper was placed between the digits to ensure no electrical contact between digits / electrode pairs. Behavioural responses required the subject to rest their dominant or the more physically agile foot on the foot pedal. For CRPS patients, this was always a foot unaffected by CRPS symptoms. To familiarise the participant with the setup and providing responses, they were given a 30s test-run of the main TSOT experiment.

The TSOT consisted of a three-way repeated measures design (Fig. 1), including the factors Change Probability (CP, levels: 10%, 30% and 50%), Side Stimulated (SS, levels: left, right hand; although for data analysis these were translated to “affected” and “unaffected” by CRPS – see below) and Change Distance (CD, levels: 1 digit, 3 digits). The experiment lasted ∼16 minutes and consisted of 890 trials, with an SOA of 1 s. One digit per trial was stimulated. On the next trial, either the same digit was stimulated (standard trial) or a different digit was stimulated (oddball trial). The probability of an oddball varied in different “Change Probability” blocks through the experiment (see Fig. 1 for block structure) such that oddballs could occur with either 10%, 30% or 50% probability. In addition, each digit was paired with another digit so that when the digit stimulated was changed, they almost always changed to the paired digit. This stimulation occurring between pairs of digits was also blocked. The four types of “Change Distance” blocks consisted of the following two-digit pairs on each of the two hands: middle to ring finger (a one-digit change, “CD1”, or adjacent pair); index to little finger (three-digit change, “CD3”). The trial sequence was randomly generated, and the same sequence used for all participants to allow comparable parameter estimates for computational modelling.

**Figure 1:**
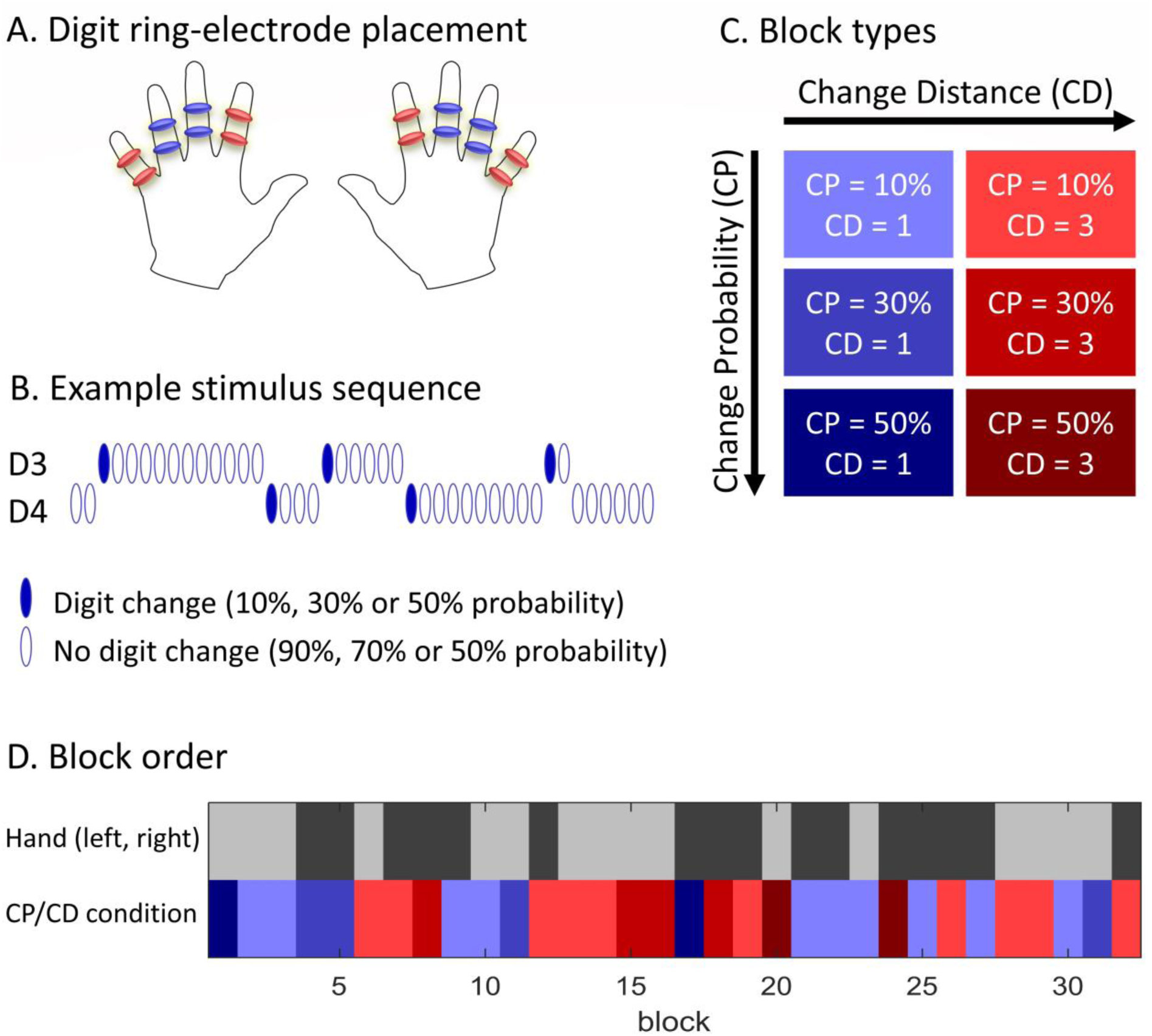
Experimental task and design: Tactile Spatial Oddball Task (TSOT) A. The TSOT stimulating digit ring electrode placement. Four electrode pairs per hand consisted of two pairs (D2 and D5) for the Change Distance 3 (CD3) condition and a further two pairs (D3 and D4) for the Change Distance 1 (CD1) condition. B. Stimuli randomly changed between the respective digits within each condition, e.g. between digits 3 and 4 (D3/4) for the CD1 condition, but with a fixed probability. Each change resulted in an “oddball” trial. C. The Change Probability (CP) constituted an orthogonal factor the Change Distance (CD) factor to provide 6 distinct block types. D. The six block types were randomised over the course of the experiment (same order for each participant to enable computational modelling). The hand stimulated was also randomly changed with an equal balance of conditions on each hand. Each block contained 30 trials. The total number of oddballs per block varied (3, 9, or 15) according to the change probability (10%, 30% or 50% respectively), as did the number of occurrences of each block type to ensure an approximately equal number of oddballs per condition across the experiment.

Participant responses were made by foot pedal release, i.e. they maintained light pressure on the foot pedal throughout but released as quickly as possible after they detected that the electrical stimulation had just shifted from one digit to another, replacing their foot back on the pedal as quickly as possible afterwards. Participants used their right foot (consistent with all being right-handed) except for the five patients whose CRPS-affected leg was on the right. In some blocks (namely, 50% oddball conditions), changes occurred rapidly, and so participants were informed beforehand they may find it difficult to keep up, but to try their best. The experiment automatically paused halfway through to give an opportunity for the participant to take a break if needed.

### Definition of “Side” conditions for analysis

During data acquisition, the ‘Side’ factor referred to stimulation of either the left or the right hand. For purpose of data analysis, the levels of the Side factor were affected and unaffected based on CRPS clinical assessment (Supplementary Table 2). This resulted in a different mixture of left and right-stimulations in each of the affected and unaffected conditions, because 10 patients were affected on the right side (5 upper limb, 5 lower limb) and 12 patients on the left side (8 upper limb, 4 lower limb). To provide adequate control for group comparisons, data in the HC group was also re-assigned as follows: to match the left/right ratio of side affected in the CRPS group, 10 healthy controls’ (randomly allocated by algorithm) right arm data and the remainder of the healthy controls’ left arm data were assigned to the ‘affected’ condition for the HC group. To be explicit, the term ‘affected’ in the HC group denotes a control condition for the CRPS ‘affected’ condition and does not imply the presence of CRPS symptoms in the HC group. The same strategy was used in a related study (Kuttikat et al., 2018).

### Behavioural data analysis

Behavioural responses were included for analysis for within-block digit-changes (i.e. within each CD1 and CD3 block type) but excluded for between-block changes (i.e. transitions between CD1 and CD3 blocks, or between hands) as there were too few of these to constitute a condition of interest for analysis. In order to assign the trial to which each response belonged, we used the following rules: (1) A minimum delay of 200ms between stimulus and response; hence responses occurring between 0 – 200ms after the stimulus were assumed to be in response to a previous trial. (2) The maximum delay between stimulus and response would be 2s. For example, if the stimulus changed on trial t and not on trial t+1, but if the response occurred on trial t+1, it was considered as occurring in response to trial t. If there was a stimulus change on trial t as well as on trial t+1 (and the response occurred at least 200ms after the t+1 stimulus) then the response was assigned to trial t+1. For trial quality assurance, we investigated whether there were group differences (CRPS vs. HC) in the proportion of targets (stimulus change trials) with missed responses (i.e. not occurring in the 200ms to 1000ms time window) and in the number of responses that were corrected according to the above rules. No statistically significant group differences were found for any condition (see Supplementary Materials for full results).

The resulting response times (RTs) were not normally distributed and so they were transformed using the natural logarithm (log-RT). Behavioural data were initially analysed using IBM SPSS software version 21 (IBM, 2012). Data that could be transformed to approximate normality were analysed using mixed ANOVA; otherwise data were analysed using non-parametric tests to investigate group differences (Mann-Whitney U test) and within-subject condition effects (Wilcoxon signed rank tests). Effect sizes are reported for non-parametric statistics using the formula *Z/√N*, where *Z* is the *z*-value output of the test and *N* is the number of samples in the test. This provides an effect size equivalent to the Pearson’s coefficient *r*, commonly interpreted as a small, medium and large effect with values of 0.1, 0.3 and 0.5 respectively.

### Data modelling: HGF model of perceptual inference

An overview of the data modelling steps for the behavioural data is illustrated in Fig. 2a, and a graphical model of the hierarchical Gaussian filter (HGF) model (Mathys et al., 2014) is shown in Fig. 2b. A detailed rationale for use of the HGF and some details of its mathematical implementation are provided in Supplementary Materials. In short, the HGF is a model of the brain’s system of hidden states. Namely, states in this case are a combination of (a) internal representations of the statistical structure of the environment, and (b) update rules that govern how internal representations change in response to sensory inputs/changes. Ultimately, the HGF acts as a model of how the brain might perceive sensory changes that have occurred. Subject-specific parameters of the model, and trajectories of hidden states in the model as they evolve over time, can be inferred from observable data.

**Figure 2:**
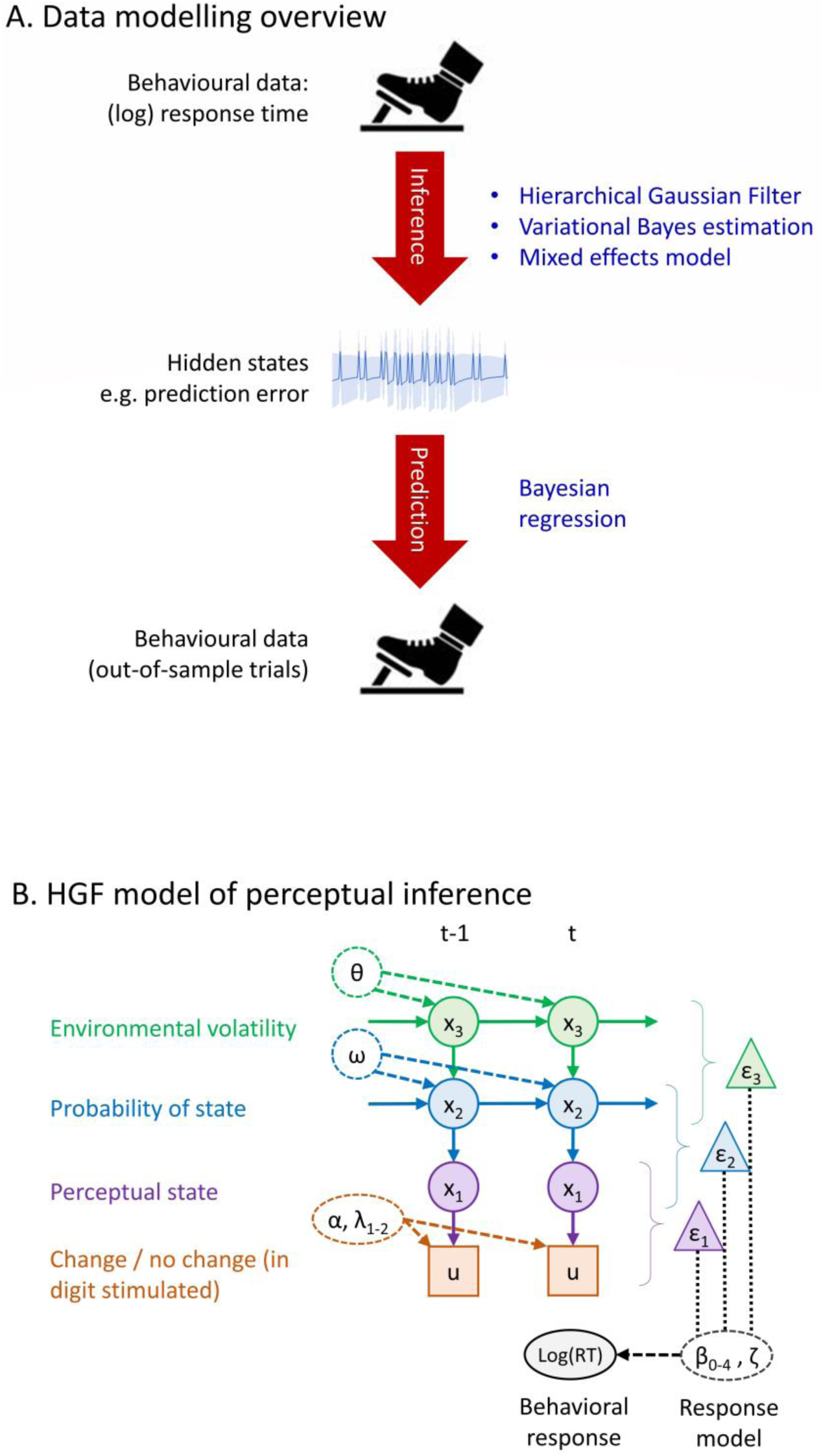
Data modelling. A. Data modelling overview. Response times (RTs) from the TSOT were used to estimate participant-specific parameters of the Hierarchical Gaussian Filter (HGF) as a model of predictive coding, resulting in the inference of internal states including prediction errors (PEs). After the best HGF model had been selected based on predictive validity (see Supplementary Materials), variational Bayes optimised the HGF parameters as part of a mixed-effects estimation model in which each participant’s HGF parameters were conditioned on empirical priors, namely the sufficient statistics of their group’s parameters (CRPS or HC group). Estimates of PEs were used to validate the model by simulating behaviour to see if the model reproduced within and between-subject differences in RTs. *B.* HGF model of perception inference. The perceptual model is comprised of three hierarchical states (x_1_, x_2_, and x_3_) that evolve over time (t), as well as binary inputs to the system (u). Solid arrows indicate conditional probabilistic relationships of a generative model of the environment (i.e. the brain’s model of how observed digit changes are generated from environmental contingencies). Inversion of the model using variational Bayes from individual log-RT data results in the estimation of participant-specific free parameters (symbols in dashed ovals) that parametrize (dashed arrows) respective inputs, outputs and hidden states: α is the sensory imprecision that is modified by λ*1* and λ*2* depending on the experimental condition; ω and θ increase the imprecision of the participants’ estimates of probability (*x_2_*) and environmental volatility (*x_3_*) respectively. The response model (grey/black) predicts log-RTs from variables in the perceptual model – in this example, from precision-weighted PEs (triangles) estimated at each level of the model – parametrized by beta weights (β). For details, see Supplementary Materials.

Two types of data were used as inputs to the HGF model: stimulus inputs (determined by the experiment) and participant responses, namely log-RTs on oddball (digit change) trials. We used RTs rather than task errors because our previous work that identified CRPS patients’ performance on a digit identification task is more clearly distinguished from control groups based on RTs (Kuttikat et al., 2018, 2016b). Stimulus inputs were coded to be binary such that each oddball (digit change trial) was 1 and each standard (no change trial) was 0. Model inversion (parameter and state estimation) from experimental data involved calculating maximum-a-posteriori (MAP) estimates for the parameters (see (Mathys et al., 2014) for details of the MAP equation and optimisation method used in the HGF toolbox).

The HGF is a modular modelling framework that allows variable model designs to be implemented and tested against data. A range of HGF models were fitted to log-RT data from the TSOT. All models were designed to differentiate theoretically important sensory and cognitive components of perceptual inference as described according to a predictive coding framework, namely prediction errors and their precision weights, which serve to update posterior estimates (on a trial-by-trial basis) of the probability of a change in the location of the stimulus.

Each model consisted of a perceptual model (the participants’ putative generative model of the environment) and response model (linking the perceptual model to participant behaviour) – detailed in Supplementary Materials. The perceptual model variations had mostly commonalities in the structure (shown in Fig. 2b) that have been described in previously published work (Mathys et al., 2014). In particular, we used a 3-level model in which the first level (x1) is the estimated perceptual state (i.e. a model of participants’ perception of change / no change), the second level (x2) represents the participants’ estimated probability of the perceived state, and the third level (x3) is the participants’ estimate of environmental volatility (likelihood of change in the probability of perceived states). States at each level are probability distributions consisting of a mean (µ) and variance (σ). States on each trial are conditioning dependent on both prior states at the same level, and the state of the next higher level, as shown by arrows in Fig. 2b. Sensory inputs to the 1^st^ level update these states according to update rules consisting of equations that have the form of PEs (see (Mathys et al., 2014) for details). Importantly, PEs occur at each level in the hierarchy: updates of x_1_ from sensory inputs are via 1^st^ level PEs (ε_1_), updates of x_2_ from changes in x_1_ are via 2^nd^ level PEs (ε_2_) and updates of x_3_ via changes in x_2_ are via 3^rd^ level PEs (ε_3_). The subject-specific parameters consist of α_0_ (variance on sensory inputs to the 1^st^ level), and ω and θ that are respectively 2^nd^ and 3^rd^ level variance parameters on states x_2_ and x_3_.

We varied both the perceptual model and in the response model in a factorial fashion (see Supplementary Fig. 1a), such that every perceptual model variation was tested with every response model variation, resulting in “families” of models with different features from the perceptual or response models. There were 3 perceptual models and 6 response models; hence all combinations of perceptual and response models resulted in 18 models. The three variations of perceptual models differed according to the number of parameters describing irreducible uncertainty (sensory noise). The six response models differed according to hypothesised components of the model driving longer RTs, which broadly categorise into either trial-by-trial estimates of prediction error, or trial-by-trial estimates of posterior uncertainty, at multiple levels in the model. We used formal model comparison methods to identify which specific variation of the HGF to take forward for comparisons between CRPS and HC groups. For purposes of model selection, data from both groups (CRPS and HC) was used jointly, such that parameters from the same model could be compared between groups. The “winning” model was taken forward for mixed-effect analyses.

### Hierarchical mixed-effect model

Optimization methods for parameter estimation are prone to errors (for example, variational Bayesian schemes can get stuck in local minima), as well as being prone to estimation error due to small sample sizes and poor parameter identifiability (Gershman, 2016). The question of group differences in model parameters is naturally framed in terms of hierarchical mixed-effect models, which condition each individual participant’s parameters on the sufficient statistics (mean and variance) of the distribution of parameters from their group, thereby providing “empirical priors” on the individual participant estimates. This has the effect of regularising individual estimates according to group statistics, which produces better individual estimates and therefore more reliable group-level tests, as well as providing better predictions of out-of-sample data, compared to parameter estimates based on fitting each individual separately (Gershman, 2016; Huys et al., 2012, 2011; Wiecki et al., 2013). The shrinkage of individual parameter estimates towards group statistics will be in proportion to how confident we are in the parameter estimates (Friston et al., 2016), i.e. subjects with uninformative data will be informed by subjects with informative data.

On this basis, the winning model (in terms of predicting RTs – previous section) was re-estimated employing a hierarchical model with empirical priors, estimated for each group separately (CRPS or HC groups). The estimation procedure is described elsewhere (Daunizeau, 2017); in sum, it involves estimating all participant’s individual model parameter a number of times (iterations), each time updating the empirical priors calculated from their group sufficient statistics from the previous iteration. As well as improving individual participants’ parameter estimates, the results also maximise the summed log-model evidence over participants. This iterative scheme was terminated when the change in free energy for both groups differed from the previous iteration by less than 1%, resulting in 7 iterations. Parameter estimates were then compared between groups using conventional non-parametric statistics (Mann-Whitney U-test). We also investigated group differences in state trajectories of interest (namely, precision-weighted PEs) using mean values over conditions in the design.

## Data availability

The data that support the findings of this study are openly available in the Open Science Framework at https://osf.io/3wh7q/.

## RESULTS

### Change probability influences behaviour more greatly under low signal-to-noise

The tactile spatial oddball task (Fig. 1c, 1d), in which participants responded whenever stimulation was switched or changed from one digit to another, provided RTs on the spatial oddball trials that participants correctly responded to. Prior to computational modelling, we initially validated the task by demonstrating modulation of RTs by orthogonal experimental conditions influencing both top-down (spatial change probability) and bottom-up (spatial change distance) factors.

Log-RTs from the task were subjected to conventional inferential statistical analysis to identify condition and group effects. A mixed ANOVA (see Supplementary Table 7 for complete statistics) revealed within-subject effects indicating that log-RTs (as shown in Fig. 3a) were sensitive to signal-to-noise of spatial changes (Change Distance, CD, *p*<0.001) and change probability (CP, *p*=0.046). Specifically, RTs were longer for smaller CDs (greater signal-to-noise) and lower CPs (more rare changes). There was also an interaction between these factors (*p*=0.001) resulting from CP effects (namely, longer RTs for rarer spatial changes) only occurring when the spatial change was small (CD1 condition). In other words, the probability of spatial change only affected behaviour when the magnitude (distance) of displacement was small. This is consistent with the notion of change detection as Bayesian inference, in which the sensory evidence (a change “signal”) is down-weighted (relative to the learnt prior expectation of a change) when there is lower signal-to-noise. Successful demonstration of these interactions supports the use of the task for generating data for purposes of fitting a computational model that integrates top-down and bottom-up influences on perception.

**Figure 3:**
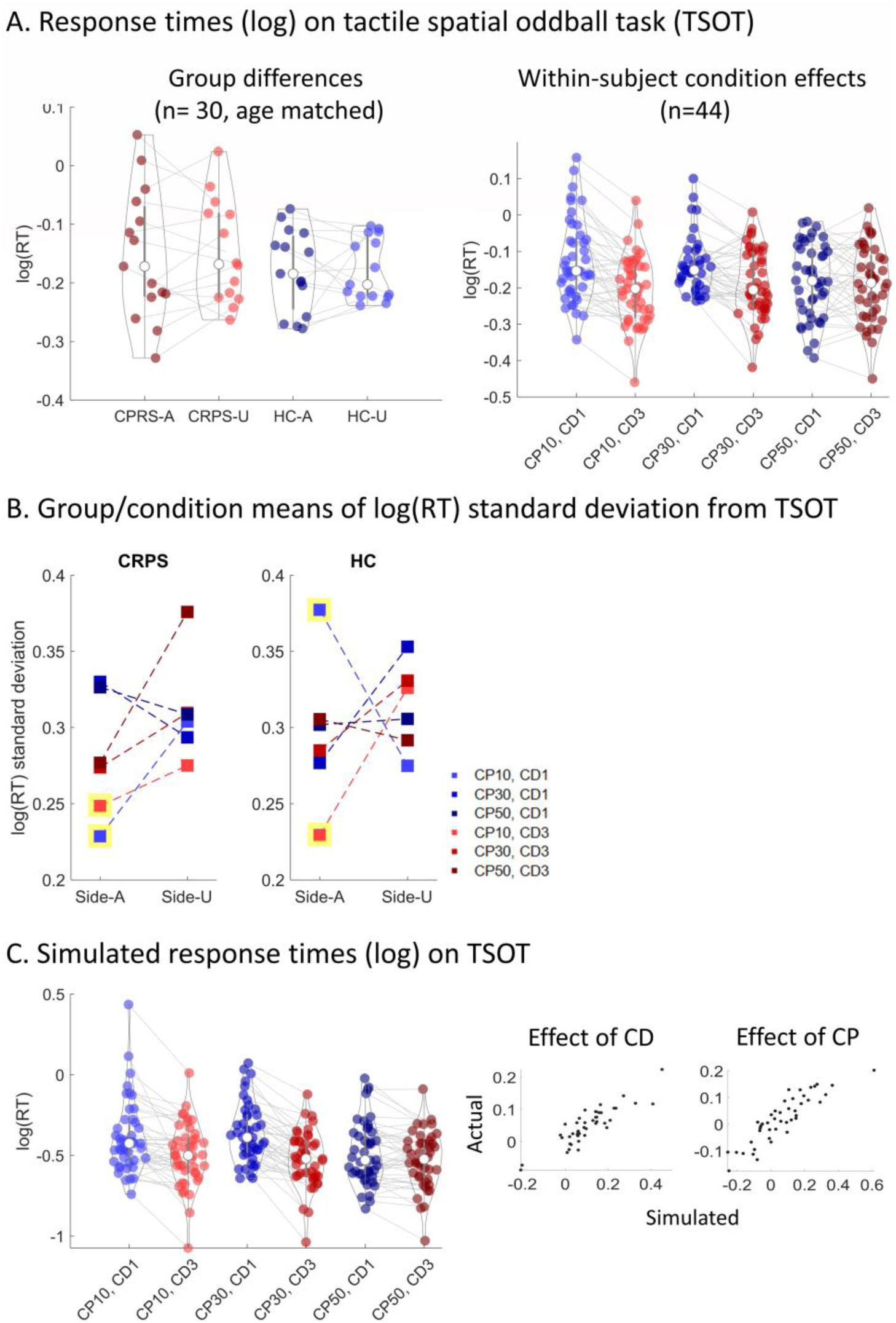
Behavioural data: descriptive plots. A. Behavioural data (response time) from TSOT task. Left: Group differences, summarised by side of stimulation (CRPS: affected vs. unaffected side, HC: pseudo-affected vs. pseudo-unaffected side – see Methods for details) and averaged over other factors in the design (Change Distance (CD) and Change Probability (CP) – see Fig. 1). P values shown are from the paired comparison of CRPS-A (affected side) vs. HC-A (pseudo-affected side) using Mann-Whitney U-tests. Right: Within-subject (condition) effects on the TSOT response times; each violin is each condition after crossing the two factors of Change Probability and Change Distance; data is averaged over the side stimulated and pooled over groups (CRPS and HC). B. Plots on the group and condition means of log(RT) standard deviation. Standard deviation was first calculated over trials, for each individual separately, to represent the variability in individual performance (RT). In this chart, data points are the mean of these SD values over all individuals in each group, for each condition. Data points outlined in yellow are the critical conditions driving a 4-way interaction effect between Group (CRPS, HC), Side (Side-A: “affected”, Side-U: “unaffected”), Change Probability condition (CP 10%, 30%, 50%) and Change Distance condition (DC 1, 3). See Results section for details. Error bars not shown for visual clarity. C. Pseudo response time data simulated by a chosen HGF model variant (perceptual model 3, response model 4 – details in Supplementary Materials) that was selected based on predictive validity, i.e. its ability to generate simulated RTs that predict actual RT data over trials of the experiment. Violin plots show the same pattern of RTs over conditions as shown in (A), while scatter plots demonstrate accurate prediction of between-participant variability in condition effects (Change Probability and Change Distance) on RT. The simulations for all HGF variants are in Supplementary Fig. 2. Violin plots show the kernel density estimate of the data (grey outlines) overlaid with means of data from each subject/condition. Also overlaid are the boxplots (median: white circle, box (IQR): thick line, whiskers (1.5x IQR): thin line). Each grey line connecting scatter-points is a subject.

### Different pattern of response time variance in CRPS patients

Group differences in *mean* log-RT for each condition were absent for the main effect of group and were also absent for all interactions involving the group factor (Supplementary Table 7), suggesting no specific or global impairments of RT in patients with CRPS compared to healthy controls. We also investigated within-subject *variance* in RTs (specifically, the standard deviation of log(RT) values (“log(RT)-SD”) over all responses within each condition) in order to identify group differences in these values. Group effects absent for the main effect of group and for two-way and three-way interactions with CD and CP factors. However, there was a 4-way interaction involving group, CP, CD and side affected (*p*=0.004, Supplementary Table 7).

Plots on the group and condition means of log(RT)-SD values (Fig. 3b) show that the significant interaction effect was driven by group differences in the CD1 vs. CD3 condition, which only occurred in the HC group when stimuli were rare (CP10: 10% change probability) and on the “affected” side. This was confirmed by follow-up tests on every CD1 vs. CD3 pair (for each Group, Side and CP combination), which found a statistically significant difference (p=0.001 uncorrected) only in the HC group, “affected” side (which is the right arm for 10/15 participants) and in the CP10 condition. No other pairs reached significance. Related to this finding, a further follow-up test confirmed a group difference (smaller in CRPS vs. HC, p=0.009 uncorrected) only on the “affected” side for the CP10 / CD1 condition, but not for any other condition. This means that for CRPS patients compared to controls, there was less variance in RTs on the affected side when there were fewer events to learn from (CP rare) and when events were not as easily distinguishable (CD short).

In order to understand the clinical relevance of this condition-specific decrease in log(RT)-SD in CRPS patients, we conducted Pearson’s correlations, within the CRPS group (n=22), between log(RT)-SD from the affected side / CP10 / CD1 condition as identified above, and four clinical variables relevant to CRPS. We found statistically significant negative relationships (consistent with the above-identified group-mean decreases in log(RT)-SD in CRPS patients compared to controls) for BPI pain severity index (r=-0.45, p=0.035) and BPI pain interference index (r=-0.63, p=0.002), but no evidence of a relationship with L/UEFI limb functioning (r=0.38, p=0.125) and NLSQ neglect-like symptoms (r=-0.35, p=0.107). Only the correlation showing more pain interference with lower log(RT)-SD values survived correction for multiple comparisons over the four clinical variables. Using hierarchical linear regression, we found the smaller log(RT)-SD from the affected side / CP10 / CD1 condition still predicted greater BPI pain interference (p=0.005) even after controlling for log(RT)-SD values from all other conditions, indicating that the relationship is specific to this condition.

### Computational modelling: Model selection and validation

Decreases in RT variance for rare and difficult to distinguish digits changes in patients with CRPS may indicate a deficit in tactile learning. To better understand these results, we employed a more sophisticated generative model of “hidden” cognitive processes theorised to modulate RTs. Computational modelling of RTs employed the hierarchical Gaussian filter (HGF, Fig. 2b; described in Methods and Supplementary Materials), a Bayesian model of predictive coding suitable for modelling volatile environments, in order to estimate prediction and PE trajectories (at 3 hierarchical levels in the model) over the course of the experiment (shown as group means in Fig. 4a,b).

**Figure 4:**
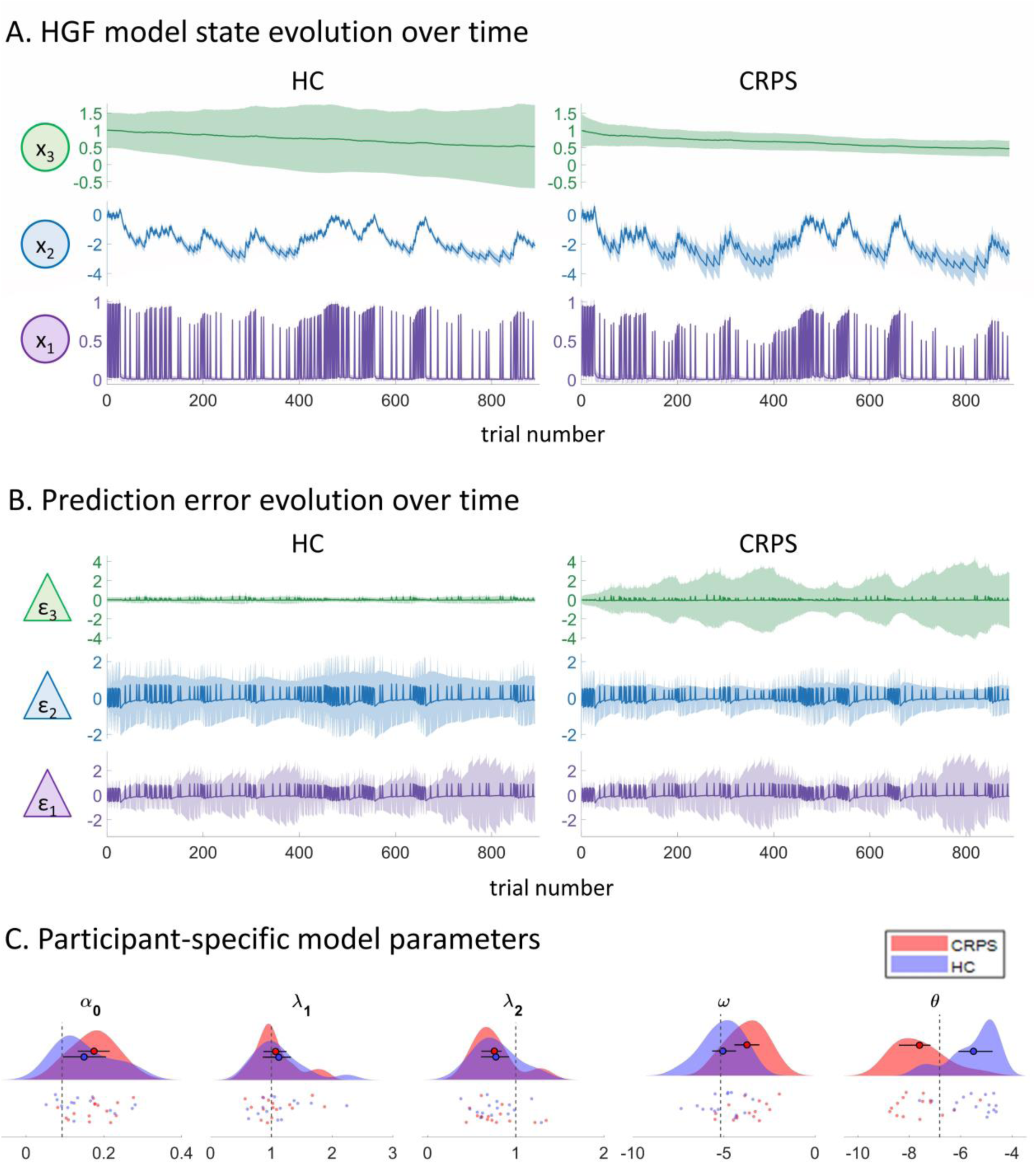
Estimated HGF model parameters and state evolution trajectories, summarised by group. A. Model state evolution over time, for each group (CRPS, HC) and each level of the HGF (x_1_, x_2_, and x_3_), resulting from fitting the HGF model to behavioural data (log-RTs) in each group. The state at each level is depicted by the mean of the distribution of each state (µ, line) and the variance (σ, shaded areas). For both mean and variance estimates, the group’s mean of these values are plotted. B. Prediction errors (PEs) and their variances over time, for each group (CRPS, HC) and each level of the HGF. The mean of the group’s PEs at each level (δ_1_, δ_2_, and δ_3_, lines) and their variances (ψ_1_, ψ_2_, and ψ_3_, shaded areas). C. Group summaries of participant-specific parameter estimates. Raincloud plots show the kernel density estimate for each group (“cloud”) and the individual participants’ estimates (“rain”), overlaid with the boxplot (circle: mean, whiskers: inter-quartile range). Vertical dashed lines show the Bayes-optimal parameter values used as priors for model fitting to each subject; comparison to group estimates indicates how far each group deviates from statistical optimality.

Model validation (see Supplementary Materials) proceeded by first identifying the best model from a factorial combination of a range of perceptual and response models that might explain trial-wise RTs. Specifically, it was important to demonstrate that the best model could simulate (reproduce) condition-effects on behaviour, as well as individual differences in these effects. Critically, the simulations used individual participant model parameters estimated from a “training set” of experimental trials, while model simulations were made on a “test set” of experimental trials that were not used to train the model (split-half cross-validation). The HGF model was chosen that was best able to simulate RTs (simulated data shown in Fig. 3c). This model consisted of a perceptual model that accounts for modulation of sensory uncertainty by two bottom-up factors in the experimental design, namely the Side (hand) stimulated and also by the Change Distance, and a response model that predicts RTs using the absolute values of precision-weighted PE calculated from the perceptual model.

### Groups differ in the precision of predictions and prediction errors

The validated HGF model was then fitted using empirical priors estimated for each group separately (n=15 per group, age matched), in order to compare parameter estimates and hidden variables of interest (predictions and PEs) between groups. Such a hierarchical estimation of model parameters has been recommended to provide more stable parameter estimates (Gershman, 2016).

There were group differences in precision-weighted PEs that indicated less efficient predictive coding of tactile-spatial change detection (Supplementary Table 9 and Fig. 4c). CRPS had larger mean values (over all conditions) of precision-weighted PEs at the 2^nd^ level (ε_2_, so-called “volatility” [i.e. variance] PE, *p*=0.007) (for further information about the difference between “value” and “volatility” PEs, see (Mathys et al., 2014)). This was driven by precision-weights at this level (ψ_2_) which showed a group difference in the same direction (*p*=0.002); conversely, unweighted PE at the 2^nd^ level (δ2) did not differ between groups (*p*=0.115). There was also a group difference in PE precision at level 3 in the HGF model (ψ_3_) with lower precision in CRPS vs. HC participants (*p*<0.001), although this did not translate to a group difference in precision-weighted PE at this level (ε_3_*, p*=0.051).

Group differences in precision weights (ψ_2_, ψ_3_) on PE were driven by subject-specific uncertainty (*inverse* precision, or imprecision) parameters, which also differed between groups. There were also corresponding differences in the variances of posterior predictions (σ_1_, σ_2_, σ_3_). Parameters differing between groups were ω and θ, which parameterize the posterior variance at the (respectively) 2^nd^ and 3^rd^ levels in the HGF (Fig. 2b and Supplementary Table 9). CRPS patients had a larger ω value (*p*=0.003) and consequently larger 2^nd^ level posterior variance (σ_2_, p=0.002), but smaller θ values (*p*<0.001) and hence smaller 3^rd^ level posterior variance (σ_3_, *p*<0.001). In other words, the results suggest that CRPS patients had less precise predictions of tactile spatial changes (HGF level 2 variance); furthermore, this imprecision was more stable (less volatile) over time (HGF level 3 variance). Exploratory cross-correlation analyses, within the CRPS group (n=15), between these two model parameters (ω, θ) and the four clinical variables relevant to CRPS did not reveal any statistically significant relationships: BPI pain severity index (ω: r=0.01, p=0.980; θ: r=0.109, p=0.699), BPI pain interference index (ω: r=-0.02, p=0.938; θ: r=-0.44, p=102), L/UEFI limb functioning (ω: r=0.03, p=0.934; θ: r=0.15, p=0.637), and NLSQ neglect-like symptoms (ω: r=0.34, p=0.219; θ: r=0.19, p=0.506).

α_0_, λ_1_ and λ_2_ parameter values (which modify the residual error variance of the model, a.k.a. irreducible uncertainty) did not significantly vary between groups (Fig. 4c). This is of interest because bottom-up sensory noise provides a lower bound on the residual error variance and would be expected to co-vary with it. Therefore, group differences were entirely related to top-down predictions, with no evidence of a role for group differences in bottom-up signal-to-noise (α_0_) or in its dependence on CD and Side conditions (λ_1_ and λ_2_ parameters respectively).

## DISCUSSION

Behavioural studies in CRPS have revealed sensory abnormalities in CRPS but underlying mechanisms remain unclear (Förderreuther et al., 2004; Lewis and Schweinhardt, 2012; Pleger et al., 2006). Here we took the novel approach of considering tactile-spatial change-detection as hierarchical Bayesian inference, as a computational model of predictive coding, to identify deficits in learning top-down predictions of tactile-spatial information. We found that CRPS patients’ predictions (of spatial displacement) were less precise, and deviate further from statistical optimality, compared to healthy controls. This imprecision was less context-dependent, such that their spatial predictions were less finely-tuned to the statistics of the environment (i.e. the temporal dynamics of sensory changes). This resulted in greater precision on prediction errors that update predictions, indicating deficits in efficient predictive coding of tactile-spatial information in patients with CRPS.

Our findings provide new insights into the possible brain mechanisms underlying abnormal tactile-spatial processing in patients with CRPS. As far we are aware, this is the first study to apply a generative mechanistic model to investigate tactile perception in chronic pain disorders. We modelled trial-by-trial dynamics in RTs to investigate statistical learning over time, providing an advance on prior research that has only considered averaged data over trials (which assumes, incorrectly, that events/responses are homogeneous and independent). This resulted in observed deficits in patients with CRPS that were not clear from condition-averaged data (mean RTs or task errors) and but were initially suggested by analysis of the variance in RTs within each condition: We found less variance in RTs on the affected side in CRPS patients (compared to controls) but this was specific to when spatial changes were rare and of a small distance. Furthermore, this reduced variance correlated with patient self-reports of interference in their daily activities from their pain symptoms.

In order to identify whether this reduced RT variance indicates a deficit in tactile learning, we tested for group differences after fitting trial-by-trial RTs to a computational model of the brain’s hidden states (e.g. predictions and PEs). By estimating specific quantities of the model and identifying how they differ between individuals, we obtain evidence for how CRPS patients learn to minimise prediction error when detecting spatial changes in tactile stimuli. Imprecision of predictions, found in CRPS patients, is calculated from two quantities in the model: firstly, the subject-specific model parameter ω, which remains static over trials, and secondly the time-varying quantity x_3_, which is updated trial-by-trial. Subject-specific parameters such as ω reflect enduring traits that are invariant to environmental changes occurring during the time-course of the experiment, but which may undergo changes over longer time-scales (possibly reflecting neural plasticity over the life-span (Bogacz, 2017)). According to the free-energy formulation of predictive coding, parameter updating (long-term plasticity) only occurs if free-energy cannot be minimised through changes in (short-term) neural activity, i.e. via the time-varying dynamics of prediction. Within the HGF model, these dynamics are modelled as the second contributor to the imprecision of predictions, namely the time-varying quantity x_3_, which is updated trial-by-trial. Updates to x_3_ are smaller when x_3_ is more precise (determined by the subject-specific parameter θ). These relationships in the model provide insights into group differences: CRPS patients had larger values of ω but smaller values of θ. This indicates that CRPS patients have more imprecise predictions but more precise estimates of environmental volatility (i.e. variability in change probabilities across the task). The net result is that the more imprecise predictions in CRPS patients are more enduring, i.e. they are not dynamically updated as greatly during the time-course of the experiment in the CRPS group. Given that parameter updating is theorised to only occur if free-energy cannot be minimised alternatively through time-varying predictions (e.g. x_3_), this implies that changes in the parameter ω in CRPS patients might be a result of deficits in learning the probabilistic structure of the task.

A corollary of greater imprecision of predictions in the CRPS group was relatively greater precision of volatility PEs (i.e. those PEs that update the participants’ putative internal model of the probabilistic structure of the environment). Greater precision of PEs results in an increased learning rate, namely greater updating of predictions in the direction indicated by the error. In other words, CRPS patients update their predictions of spatial changes at least as well as healthy controls, but those resulting predictions are less precise and less finely-tuned to statistics of the environment. This means that tactile-spatial change detection in patients with CRPS is less influenced by the history of sensory changes, and more driven by current or very recent sensory inputs.

The results provide a more nuanced view of somatosensory pathology in CRPS as commonly hypothesised in the literature, emphasising the role of higher-order learning rather than low-level sensory representations. We found that, despite more imprecise predictions of tactile-spatial changes, there were no group differences in model error variances (“irreducible uncertainty” (de Berker et al., 2016)), nor the additional model parameters that describe how this low-level uncertainty varies across digit change distances and the hand stimulated. This provides evidence against the view that sensory signal-to-noise, which would be expected to decrease this low-level sensory uncertainty, is deficient in CRPS. Our findings are consistent with a nascent body of evidence against the view of primary somatosensory dysfunction in CRPS pathology as suggested by recent MRI findings of a lack of S1 differences in the somatotopic representation of the affected limb (Di Pietro et al., 2015; Mancini et al., 2018; van Velzen et al., 2016). There is preliminary evidence that anterior insula cortex (ACC), which codes both tactile and pain-related PEs (Allen et al., 2015; Seymour et al., 2004), exhibits abnormal functional and structural (white-matter) connections in the brains of patients with CRPS (Geha et al., 2008; Marinus et al., 2011). A closely connected structure, the ACC, can be considered a hierarchically high-level hub in sensory perception, sub-serving a broad range of functions and integrating emotional and autonomic responses (Craig, 2003) to sensory information. Our findings are more consistent with abnormalities in ACC and other regions in CRPS that are involved in hierarchically higher levels of sensory learning, possibly involving working memory (e.g. see Cashdollar et al., 2017).

Two key strengths of our analysis were to firstly validate the tactile-spatial change-detection task and secondly to validate the computation model, which are important steps prior to attempting to infer differences between CRPS and healthy controls. This was achieved by statistically predicting behaviour (RTs) from factors in the experimental design (validating the task) and from simulations of the computational model (validating the model). Specifically, RTs were modulated by both top-down (spatial change probability) and bottom-up (spatial change distance) factors, in a manner consistent with their putative role as markers of precision-weighted PEs in the brain (Kuttikat et al., 2018, 2016a). Subsequently, the computational model was able to simulate (reproduce) these condition-effects on behaviour and individual differences in these condition effects. These validation steps provided the foundation for investigating group differences in model parameters.

Several limitations of our approach merit further research. First, it is unknown whether the model parameter estimates are stable over time (test-retest reliability) and over what periods of time – for the results to be clinically meaningful, at least short-term stability (e.g. over days/weeks) would be a requirement. This also impacts on the likely success of interventions designed to improve sensory learning. Second, our results do not specify a somatosensory-specific deficit in patients with CRPS; further studies can employ control tasks to test if the findings point to a more general deficit in modelling environmental statistics and contingencies.

Finally, deficits in statistical learning may have therapeutic implications. Statistical learning is important to enable individuals to extract patterns or regularities from the environment over time (Hasson, 2017), and by extension, make predictions that facilitate adaptive behaviour. Because the world is both uncertain and changing, when a sensory change occurs, what has been learned must be revised: learning should therefore be flexible (Heilbron and Meyniel, 2019). Our results suggest that patients with CRPS utilise a computational strategy of only taking into account recent observations (and forgetting about the remote past), which has been theorised to be a computationally cheaper way of enabling flexibility in learning compared to engaging in hierarchically higher-level learning of probabilistic structure (Heilbron and Meyniel, 2019). Such a strategy may be compensatory or adaptive for simple tasks but may be disadvantageous when complex information needs to be learnt over longer periods. This may have implications for therapies based on sensory re-training whose effects may be limited in extent or short-lived if success depends on higher-order learning. For example, sensory discrimination training, which has some supportive evidence in its favour (Moseley et al., 2008), aims to reduce tactile discrimination thresholds as a marker of improved spatial precision in somatosensory processing. We might speculate that patients can improve at simple spatial discrimination tasks (possibly providing some initial therapeutic benefit) while still experiencing difficulty in higher-order learning. Insofar as such high-order learning turns out to be important for CRPS pathology, improvement in simple sensory discrimination may either provide limited or short-lived therapeutic benefit.

## ACKNOWLEDGEMENTS

The authors would like to thank the participants in this study. We acknowledge the support from the NIHR Cambridge Clinical Research Facility, and from the UK CRPS Registry, who facilitated identification of potential participants. The research was funded by an EFIC-Grunenthal Grant awarded to CB and a Pain Relief Foundation grant awarded to CB, NS and MCL, with additional support from the NIHR Clinical Research Network (Eastern) and Versus Arthritis for IS. The authors declare no competing interests.

## AUTHOR CONTRIBUTIONS

CB, NS and MCL substantially contributed to the conception of this work; CB and MCL designed the experiments and wrote the paper; NS, IS and MCL substantially contributed to the recruitment of participants; IS conducted the experiments. All authors gave final approval to the version of the paper to be published.

## SUPPLEMENTARY MATERIALS

### Participant recruitment and characteristics

Potential CRPS participants were identified from either local Rheumatology databases of CRPS patients at Cambridge University Hospitals (CUH), or from the national UK CRPS Registry. The total number of potentially eligible CRPS patients contacted was 65, of which 22 patients were recruited. All patients were diagnosed with unilateral upper or lower limb CRPS (ruling out CRPS on the unaffected side) according to modified Budapest Research Criteria (Harden et al., 2007). The inclusion criteria were kept as broad as possible, including upper and lower limb-affected patients on the left or the right side. Although the study tests involved tactile stimulation on the hand only, previous work (Kuttikat et al., 2018, 2016b) found that the location of CRPS symptoms (namely, upper vs. lower limb) did not significantly affect behavioural performance in discrimination tasks. Healthy controls (HCs) were recruited by advertising the study using posters in CUH. 23 HCs were recruited with the aim of matching to CRPS patients in age and sex. Data from 1 HC were excluded from the study analysis due to extremely noisy EEG data that could not be corrected or removed. This resulted in 22 in the HC group. For statistical comparisons, the groups were reduced to n=15 each matched by age. Demographic and medical details in Supplementary Tables 1 and 2.

**Supplementary Table 1:** Participant characteristics.

| Characteristic | Whole sample, n=44 |  |  |  | Age matched, n=30 |  |  |  |
| --- | --- | --- | --- | --- | --- | --- | --- | --- |
|  | CRPS (n=22) | Healthy (n=22) | Statistic (Z, Chi) | p value | CRPS (n=15) | Healthy (n=15) | Statistic (Z, Chi) | p value |
| Female (%) | 17 (77.3) | 15 (68.2) | 0.46<sup>a</sup> | 0.5 | 12 (80.0) | 11 (73.3) | 0.19 | 0.666 |
| Age in years, median (range) | 57 (20-74) | 40.5 (19-61) | 2.22<sup>b</sup> | 0.026 | 50 (20-61) | 49 (19-61) | 0.08 | 0.934 |
| DIT, affected side, % accuracy, median (range) | 10 (3-10) | 10 (7-10) | 0.38<sup>b</sup> | 0.748 | 10 (3-10) | 10 (7-10) | 0.03 | 0.98 |
| DIT, unaffected side, % accuracy, median (range) | 10 (9-10) | 10 (6-10) | 0.52<sup>b</sup> | 0.605 | 10 (9-10) | 10 (6-10) | 0.14 | 0.888 |
| DIT, affected side, response time, median (range) | 20 (13-70) | 16.5 (13-27) | 2.45<sup>b</sup> | 0.014 | 18 (13-70) | 17 (13-27) | 1.11 | 0.269 |
| DIT, unaffected side, response time, median (range) | 18 (12-33) | 17.5 (13-26) | 0.64<sup>b</sup> | 0.523 | 18 (12-33) | 18 (14-26) | 0.08 | 0.933 |
| PCS total, median (range) | 25 (0-47) | 8.5 (0-30) | 3.02<sup>b</sup> | 0.003 | 25 (6-47) | 7 (0-30) | 3.64 | <0.001 |
| HADS anxiety, median (range) | 9 (2-16) | 3.5 (0-10) | 3.52<sup>b</sup> | <0.001 | 9 (4-16) | 5 (0-10) | 3.48 | 0.001 |
| HADS depression, median (range) | 8 (1-12) | 1 (0-7) | 5.23<sup>b</sup> | <0.001 | 10 (4-12) | 0 (0-7) | 4.60 | <0.001 |

<sup>a</sup> Chi-square test
<sup>b</sup> Mann-Whitney U test, Z score
DIT: Digit identification test; PCS: Pain catastrophizing scale; HADS: Hospital anxiety and depression scale

**Supplementary Table 2:**
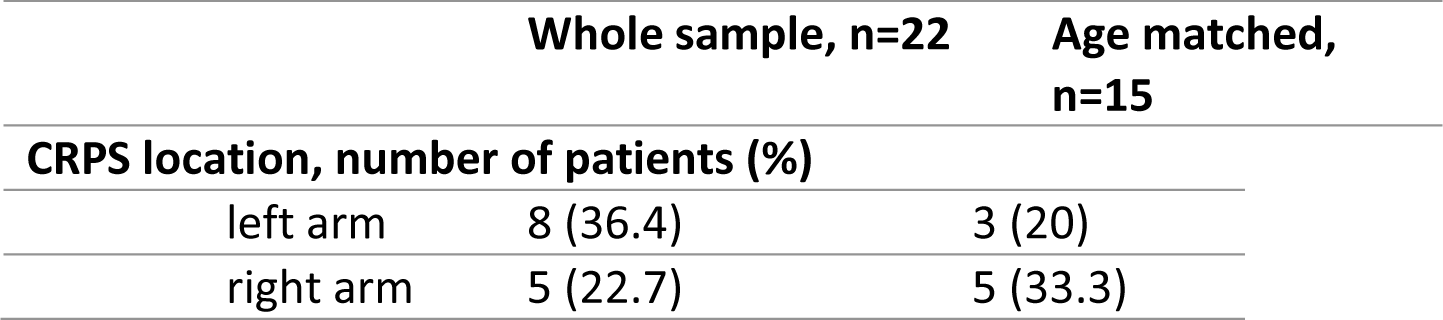
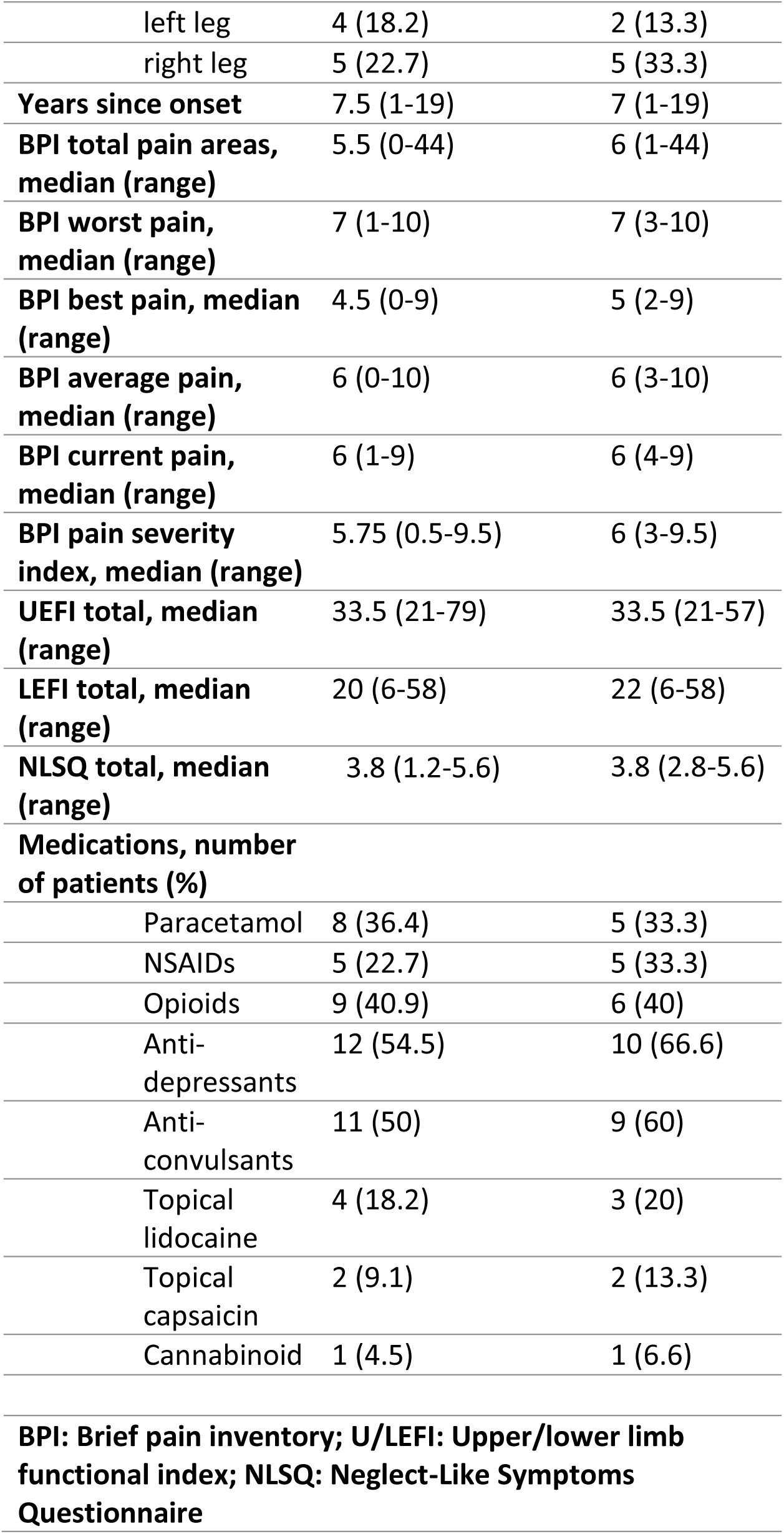
Patient medical characteristics.

|  | Whole sample, n=22 | Age matched, n=15 |
| --- | --- | --- |
| **CRPS location, number of patients (%)** |  |  |
| left arm | 8 (36.4) | 3 (20) |
| right arm | 5 (22.7) | 5 (33.3) |
| left leg | 4 (18.2) | 2 (13.3) |
| right leg | 5 (22.7) | 5 (33.3) |
| **Years since onset** | 7.5 (1-19) | 7 (1-19) |
| **BPI total pain areas, median (range)** | 5.5 (0-44) | 6 (1-44) |
| **BPI worst pain, median (range)** | 7 (1-10) | 7 (3-10) |
| **BPI best pain, median (range)** | 4.5 (0-9) | 5 (2-9) |
| **BPI average pain, median (range)** | 6 (0-10) | 6 (3-10) |
| **BPI current pain, median (range)** | 6 (1-9) | 6 (4-9) |
| **BPI pain severity index, median (range)** | 5.75 (0.5-9.5) | 6 (3-9.5) |
| **UEFI total, median (range)** | 33.5 (21-79) | 33.5 (21-57) |
| **LEFI total, median (range)** | 20 (6-58) | 22 (6-58) |
| **NLSQ total, median (range)** | 3.8 (1.2-5.6) | 3.8 (2.8-5.6) |
| **Medications, number of patients (%)** |  |  |
| Paracetamol | 8 (36.4) | 5 (33.3) |
| NSAIDs | 5 (22.7) | 5 (33.3) |
| Opioids | 9 (40.9) | 6 (40) |
| Anti-depressants | 12 (54.5) | 10 (66.6) |
| Anti-convulsants | 11 (50) | 9 (60) |
| Topical lidocaine | 4 (18.2) | 3 (20) |
| Topical capsaicin | 2 (9.1) | 2 (13.3) |
| Cannabinoid | 1 (4.5) | 1 (6.6) |
| **BPI: Brief pain inventory; U/LEFI: Upper/lower limb functional index; NLSQ: Neglect-Like Symptoms Questionnaire** |  |  |

### Participant recruitment criteria

#### Inclusion criteria

- Participant is willing and able to give informed consent for participation in the study.
- Male or Female, aged 18-80 years.
- Right handed
- Able to communicate fluently in English
- Healthy Volunteers, or
- Diagnosed with unilateral upper or lower limb Complex Regional Pain Syndrome according to modified Budapest Research Criteria given below, or
- Unilateral fracture of the upper or lower limb in the absence of any symptoms of Complex Regional Pain Syndrome.

Exclusion criteria: The participant may not enter the study if ANY of the following apply:

- Previous or current diagnosis of peripheral neuropathy, stroke, Transient Ischemic Attack, multiple sclerosis, malignancy or seizure disorder
- Unable to communicate fluently in English
- Pregnant
- Unable to or unwilling to give informed consent

Modified Budapest Research Criteria for diagnosis of CRPS (Harden et al., 2007):

1. Continuing pain, which is disproportionate to any inciting event
2. Must report at least one symptom in all *four* of the following categories:

Sensory: Reports of hyperesthesia and/or allodynia
Vasomotor: Reports of temperature asymmetry and/or skin colour changes and/or skin colour asymmetry
Sudomotor/ Oedema: Reports of oedema and/or sweating changes and/or sweating asymmetry
Motor/Trophic: Reports of decreased range of motion and/or motor dysfunction (weakness, tremor, dystonia) and/or trophic changes (hair, nail, skin)
3. Must display at least one sign at time of evaluation in *two or more* of the following categories:

Sensory: Evidence of hyperalgesia (to pinprick) and/or allodynia (to light touch and/or temperature sensation and/or deep somatic pressure and/or joint movement)
Vasomotor: Evidence of temperature asymmetry (>1°C) and/or skin colour changes and/or asymmetry
Sudomotor/Oedema: Evidence of oedema and/or sweating changes and/or sweating asymmetry
Motor/Trophic: Evidence of decreased range of motion and/or motor dysfunction (weakness, tremor, dystonia) and/or trophic changes (hair, nail, skin)
4. There is no other diagnosis that better explains the signs and symptoms.

### Sample size

A formal sample size calculation was not performed. Sample size was informed by previous research into CRPS at this institution (Kuttikat et al., 2018) which identified altered behaviour and cortical responses to tactile stimulation while using similar tasks. This was observable at statistical significance with 13 CRPS patients and 13 matched healthy controls. We surpassed the sample size by recruiting 22 CRPS patients and 23 healthy controls, so that age-matched group comparisons can be made on a subset of n=15 per group.

### Clinical evaluation

Participants with CRPS completed standardised questionnaires to assess their pain and neglect-like symptoms: Neglect-like Symptom Questionnaire (Galer and Jensen, 1999), and Brief Pain Inventory (Cleeland and Ryan, 1994). Physical functioning was assessed using the Upper Extremity Functional Index (Stratford et al., 2001) and Lower Extremity Functional Index (Binkley et al., 1999), while emotional functioning was assessed using the Hospital Anxiety and Depression Scale (Thorlund, 2010). After completing questionnaires, participants underwent a Digit Identification Task (DIT) (Förderreuther et al., 2004; Kuttikat et al., 2016b), which was developed to quantify tactile spatial disturbances in CRPS clinically.

### Digit identification task (DIT)

To find out whether our CRPS sample was qualitatively different in perceptual performance compared to our previous studies in this population (Kuttikat et al., 2018, 2016b), we tested whether CRPS patients took longer to complete a digit identification task. This task requires the more complex cognitive process of discriminating and reporting on which digit was stimulated (as opposed to the more cognitively simple TSOT). The DIT procedure involved 10 touches, in a predefined order (unknown to the participant – perceived as effectively random), to the digits of each hand in turn (one touch per digit). Touches were from the index finger of the experimenter and involve gentle but clearly perceived mild stroke of the dorsal part of the middle segment of each digit. No adjacent digit was touched in consecutive sequence.

During the task, participants had their eyes closed and were seated with left and right hands placed on their respective thighs. Participants were asked to respond to each touch verbally, calling a thumb touch number “1”, index finger “2” and so on to the little finger. The task was first completed on the affected hand (if a CRPS patient) or on a randomly allocated hand (if a healthy control). For each hand separately, the time was measured from when the first digit was touched to when the last answer was given. The number of correct and incorrect answers were recorded.

Comparing patients to controls in the whole sample (n=44), on average, patients took longer to complete the task but only on their affected side (p=0.014, Supplementary Table 1). CRPS patients were not less accurate on average, on either side. However, in age-matched group analysis (n=30), we did not find that CRPS patients took longer to complete the digit identification task, nor where they less accurate, on either affected or unaffected sides on average (Supplementary Table 1). Although 4 out of the 15 CRPS patients in the age-matched group took longer on the task than the healthy control with the longest completion time, this was not sufficient to result in a statistically significant difference between groups. The results highlight the heterogeneity of patients according to such performance metrics.

### Tactile spatial oddball task (TSOT): Electrical current delivered for each group

On average, there was no significant difference in the electrical current (in mA) delivered between CRPS patients and HC participants when considering either all n=44 participants (CRPS: M=6.87, SD=1.87; HC: M=6.54, SD=2.00; p=0.578) nor the n=30 matched subgroup (CRPS: M=6.66, SD=1.86; HC: M=6.92, SD=1.90; p=0.708). We also tested for relationships between current delivered and key dependent variables (mean RT and mean accuracy) in each group separately (n=22 per group) using Spearman correlations (two-tailed). There was a significant positive correlation (rho=0.577, p=0.005 (uncorrected)) between the current delivered and mean RTs (across all trials/conditions) in the HC group only. Since current delivered was a multiple of sensory threshold in this group, we can conclude that individuals who were less sensitive to detecting tactile stimuli also took longer to respond to changes in their location. There were no significant correlations found in the CRPS group, nor did stimulus intensity relate to computational model parameter values within either of the n=15 subgroups.

### Tactile spatial oddball task (TSOT): Quality control analysis on missed and corrected responses

For quality control purposes, we investigated whether there were group differences (n=15 per matched group) in responses to targets (digit change trials) in order to assess whether the above rules for assigning responses to targets disproportionately affected the CRPS or HC group. We calculated three metrics: (1) “missed response”: proportion of trials in which there was a digit change (target for response) but no response within the 200ms to 1000ms time window post-stimulus, (2) “corrected response”: proportion of trials in which there was a “missed response” according to the above definition but there was a response occurring later that was assigned to the current trial according to the rules described in Methods, (2) “fraction corrected”: the fraction of “corrected responses” over the total number of “missed responses”. For each of the above three metrics, we provide the mean and SD for each condition and each group, as well as the Z-scores and p values for independent non-parametric tests between the two groups (Supplementary Table 3). Comparing the means, there was a tendency for the CRPS group to have more missed responses and a greater number of corrected responses, but this was not consistent across all conditions, even for the more challenging CP-50 (50% change probability) conditions. No statistically significant group differences were found for any of the conditions on any metric.

**Supplementary Table 3:** Proportion of missed and corrected trials on Tactile Spatial Oddball Task (TSOT)

|  | Change probability: 10% |  |  |  | Change probability: 30% |  |  |  | Change probability: 50% |  |  |  |
| --- | --- | --- | --- | --- | --- | --- | --- | --- | --- | --- | --- | --- |
|  | Side: affected |  | Side: unaffected |  | Side: affected |  | Side: unaffected |  | Side: affected |  | Side: unaffected |  |
| <u>Missed responses</u> |  |  |  |  |  |  |  |  |  |  |  |  |
| Descriptive statistics | CD1 | CD3 | CD1 | CD3 | CD1 | CD3 | CD1 | CD3 | CD1 | CD3 | CD1 | CD3 |
| Group mean (CRPS) | 0.447 | 0.220 | 0.427 | 0.193 | 0.575 | 0.313 | 0.529 | 0.375 | 0.619 | 0.376 | 0.605 | 0.448 |
| Group SD (CRPS) | 0.393 | 0.276 | 0.310 | 0.175 | 0.264 | 0.254 | 0.274 | 0.230 | 0.209 | 0.304 | 0.198 | 0.170 |
| Group mean (HC) | 0.407 | 0.233 | 0.360 | 0.187 | 0.529 | 0.463 | 0.500 | 0.350 | 0.514 | 0.467 | 0.481 | 0.386 |
| Group SD (HC) | 0.225 | 0.206 | 0.275 | 0.203 | 0.192 | 0.216 | 0.274 | 0.226 | 0.216 | 0.282 | 0.194 | 0.246 |
| Mann-Whitney U test |  |  |  |  |  |  |  |  |  |  |  |  |
| Effect size: Z statistic | 0.188 | -0.530 | 0.605 | 0.511 | 0.521 | -1.624 | 0.146 | 0.376 | 1.295 | -0.855 | 1.381 | 0.709 |
| P value (asymptotic 2-tailed) | 0.851 | 0.596 | 0.545 | 0.609 | 0.603 | 0.104 | 0.884 | 0.707 | 0.195 | 0.392 | 0.167 | 0.478 |
| Corrected p value | 10.212 | 7.152 | 6.540 | 7.308 | 7.236 | 1.248 | 10.608 | 8.484 | 2.340 | 4.704 | 2.004 | 5.736 |
| <u>Corrected responses</u> |  |  |  |  |  |  |  |  |  |  |  |  |
| Descriptive statistics | CD1 | CD3 | CD1 | CD3 | CD1 | CD3 | CD1 | CD3 | CD1 | CD3 | CD1 | CD3 |
| Group mean (CRPS) | 0.147 | 0.133 | 0.220 | 0.120 | 0.108 | 0.092 | 0.100 | 0.125 | 0.086 | 0.086 | 0.110 | 0.148 |
| Group SD (CRPS) | 0.173 | 0.188 | 0.214 | 0.108 | 0.080 | 0.100 | 0.102 | 0.118 | 0.077 | 0.090 | 0.108 | 0.087 |
| Group mean (HC) | 0.180 | 0.120 | 0.140 | 0.093 | 0.092 | 0.129 | 0.063 | 0.075 | 0.076 | 0.090 | 0.100 | 0.090 |
| Group SD (HC) | 0.142 | 0.115 | 0.155 | 0.103 | 0.070 | 0.093 | 0.058 | 0.098 | 0.079 | 0.095 | 0.080 | 0.099 |
| Mann-Whitney U test |  |  |  |  |  |  |  |  |  |  |  |  |
| Effect size: Z statistic | -1.034 | 0.394 | 0.989 | 0.698 | 0.365 | -1.105 | 0.762 | 1.320 | 0.434 | -0.065 | 0.022 | 1.919 |
| P value (asymptotic 2-tailed) | 0.301 | 0.694 | 0.323 | 0.485 | 0.715 | 0.269 | 0.446 | 0.187 | 0.664 | 0.948 | 0.983 | 0.055 |
| Corrected p value | 3.612 | 8.328 | 3.876 | 5.820 | 8.580 | 3.228 | 5.352 | 2.244 | 7.968 | 11.376 | 11.796 | 0.660 |
| <u>Fraction of corrected/missed</u> |  |  |  |  |  |  |  |  |  |  |  |  |
| Descriptive statistics | CD1 | CD3 | CD1 | CD3 | CD1 | CD3 | CD1 | CD3 | CD1 | CD3 | CD1 | CD3 |
| Group mean (CRPS) | 0.485 | 0.730 | 0.501 | 0.671 | 0.200 | 0.277 | 0.211 | 0.351 | 0.134 | 0.226 | 0.190 | 0.316 |
| **Group SD (CRPS)** | 0.390 | 0.351 | 0.321 | 0.406 | 0.147 | 0.237 | 0.214 | 0.312 | 0.126 | 0.193 | 0.194 | 0.170 |
| **Group mean (HC)** | 0.465 | 0.573 | 0.325 | 0.630 | 0.191 | 0.296 | 0.175 | 0.202 | 0.145 | 0.244 | 0.191 | 0.266 |
| **Group SD (HC)** | 0.294 | 0.323 | 0.335 | 0.335 | 0.158 | 0.268 | 0.253 | 0.289 | 0.121 | 0.282 | 0.133 | 0.224 |
| **Mann-Whitney U test** |  |  |  |  |  |  |  |  |  |  |  |  |
| **Effect size: Z statistic** | 0.139 | 1.137 | 1.312 | 0.483 | 0.021 | -0.074 | 0.951 | 1.415 | -0.296 | -0.099 | -0.610 | 0.928 |
| **P value (asymptotic 2-tailed)** | 0.889 | 0.256 | 0.190 | 0.629 | 0.983 | 0.941 | 0.342 | 0.157 | 0.767 | 0.921 | 0.542 | 0.354 |
| **Corrected p value** | 10.668 | 3.072 | 2.280 | 7.548 | 11.796 | 11.292 | 4.104 | 1.884 | 9.204 | 11.052 | 6.504 | 4.248 |

### Tactile spatial oddball task (TSOT): Accuracy

Accuracy data from the TSOT (Supplementary Table 4) were not used for further modelling due to ceiling effects in the rare change-probability (10%) condition. Due to these ceiling effects the data did not approximate normality well enough for parametric statistics. Regarding within-subject effects, a 2-tailed Friedman test found a significant effect of change probability (medians: 10% condition = 98.3, 30% condition = 89.2, 50% condition = 81.0; p < 0.001), while 2-tailed Wilcoxon signed-rank tests found a significant effect of digit change distance (medians: CD1 = 86.3, CD3 = 90.5, p < 0.001) but not side stimulated (medians: affected = 88.8, unaffected = 86.3; p = 0.290). We compared groups using Mann-Whitney U tests on each cell in the design (correcting for multiple comparisons) and no statistically significant group differences were identified.

**Supplementary Table 4:**
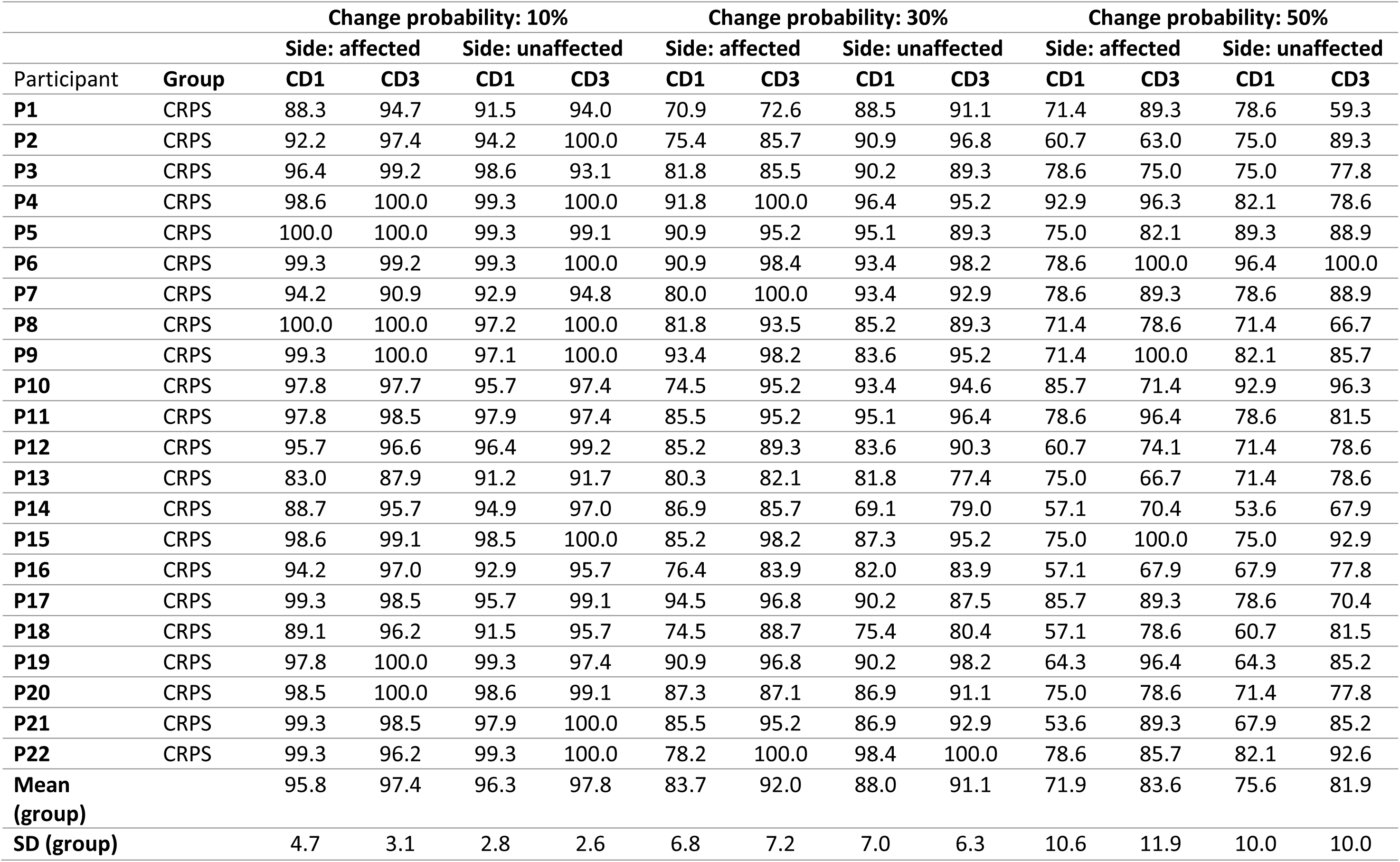
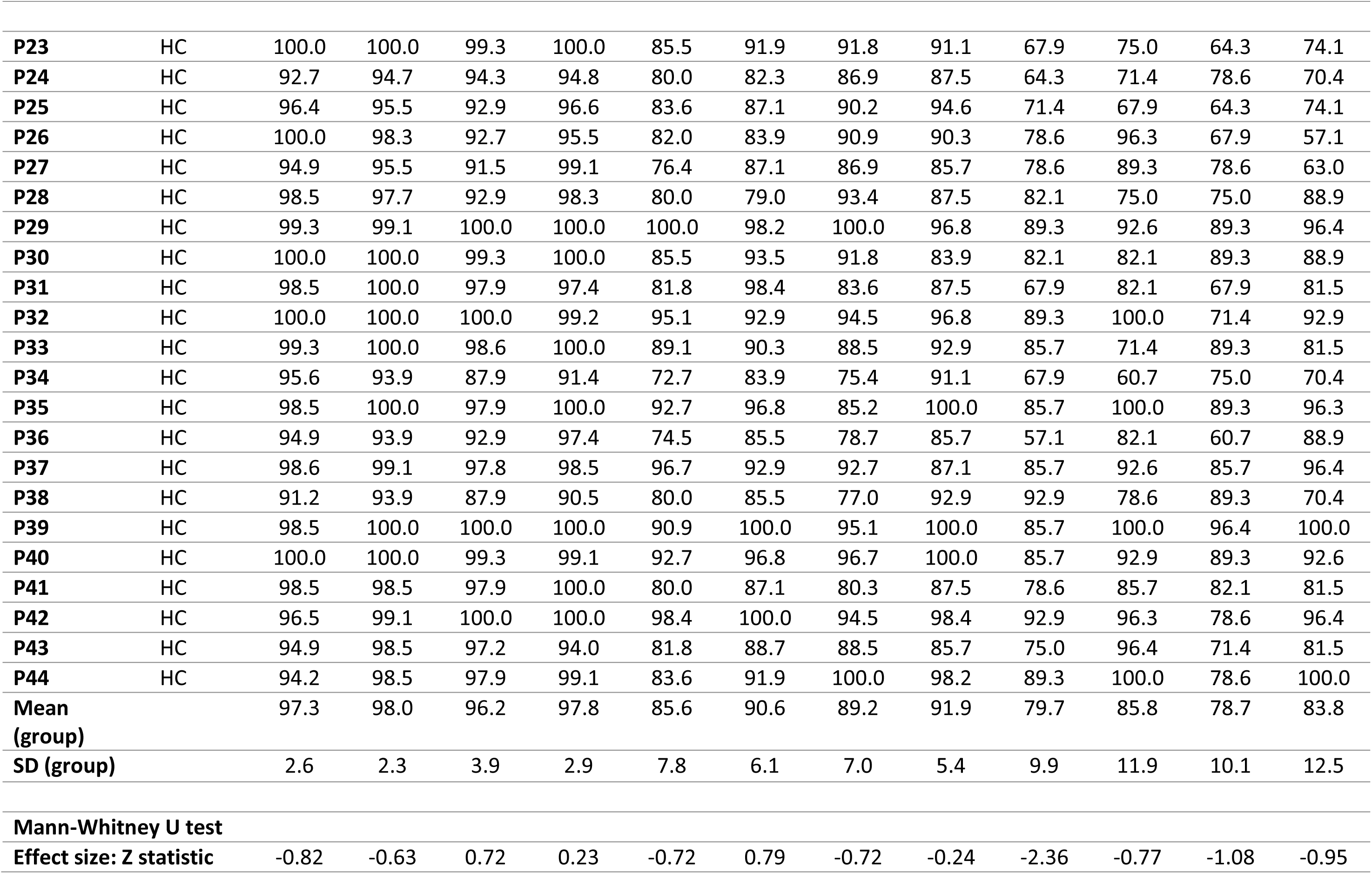
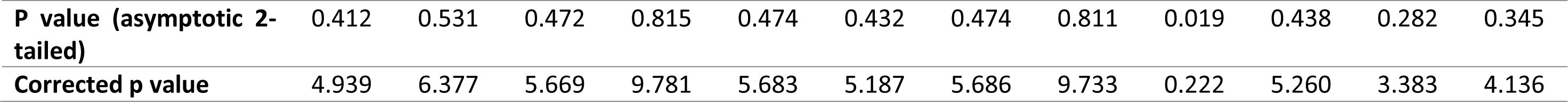
Tactile Spatial Oddball Task (TSOT) accuracy data.

| Participant | Group | Change probability: 10% |  |  |  | Change probability: 30% |  |  |  | Change probability: 50% |  |  |  |
| --- | --- | --- | --- | --- | --- | --- | --- | --- | --- | --- | --- | --- | --- |
|  |  | Side: affected |  | Side: unaffected |  | Side: affected |  | Side: unaffected |  | Side: affected |  | Side: unaffected |  |
|  |  | CD1 | CD3 | CD1 | CD3 | CD1 | CD3 | CD1 | CD3 | CD1 | CD3 | CD1 | CD3 |
| P1 | CRPS | 88.3 | 94.7 | 91.5 | 94.0 | 70.9 | 72.6 | 88.5 | 91.1 | 71.4 | 89.3 | 78.6 | 59.3 |
| P2 | CRPS | 92.2 | 97.4 | 94.2 | 100.0 | 75.4 | 85.7 | 90.9 | 96.8 | 60.7 | 63.0 | 75.0 | 89.3 |
| P3 | CRPS | 96.4 | 99.2 | 98.6 | 93.1 | 81.8 | 85.5 | 90.2 | 89.3 | 78.6 | 75.0 | 75.0 | 77.8 |
| P4 | CRPS | 98.6 | 100.0 | 99.3 | 100.0 | 91.8 | 100.0 | 96.4 | 95.2 | 92.9 | 96.3 | 82.1 | 78.6 |
| P5 | CRPS | 100.0 | 100.0 | 99.3 | 99.1 | 90.9 | 95.2 | 95.1 | 89.3 | 75.0 | 82.1 | 89.3 | 88.9 |
| P6 | CRPS | 99.3 | 99.2 | 99.3 | 100.0 | 90.9 | 98.4 | 93.4 | 98.2 | 78.6 | 100.0 | 96.4 | 100.0 |
| P7 | CRPS | 94.2 | 90.9 | 92.9 | 94.8 | 80.0 | 100.0 | 93.4 | 92.9 | 78.6 | 89.3 | 78.6 | 88.9 |
| P8 | CRPS | 100.0 | 100.0 | 97.2 | 100.0 | 81.8 | 93.5 | 85.2 | 89.3 | 71.4 | 78.6 | 71.4 | 66.7 |
| P9 | CRPS | 99.3 | 100.0 | 97.1 | 100.0 | 93.4 | 98.2 | 83.6 | 95.2 | 71.4 | 100.0 | 82.1 | 85.7 |
| P10 | CRPS | 97.8 | 97.7 | 95.7 | 97.4 | 74.5 | 95.2 | 93.4 | 94.6 | 85.7 | 71.4 | 92.9 | 96.3 |
| P11 | CRPS | 97.8 | 98.5 | 97.9 | 97.4 | 85.5 | 95.2 | 95.1 | 96.4 | 78.6 | 96.4 | 78.6 | 81.5 |
| P12 | CRPS | 95.7 | 96.6 | 96.4 | 99.2 | 85.2 | 89.3 | 83.6 | 90.3 | 60.7 | 74.1 | 71.4 | 78.6 |
| P13 | CRPS | 83.0 | 87.9 | 91.2 | 91.7 | 80.3 | 82.1 | 81.8 | 77.4 | 75.0 | 66.7 | 71.4 | 78.6 |
| P14 | CRPS | 88.7 | 95.7 | 94.9 | 97.0 | 86.9 | 85.7 | 69.1 | 79.0 | 57.1 | 70.4 | 53.6 | 67.9 |
| P15 | CRPS | 98.6 | 99.1 | 98.5 | 100.0 | 85.2 | 98.2 | 87.3 | 95.2 | 75.0 | 100.0 | 75.0 | 92.9 |
| P16 | CRPS | 94.2 | 97.0 | 92.9 | 95.7 | 76.4 | 83.9 | 82.0 | 83.9 | 57.1 | 67.9 | 67.9 | 77.8 |
| P17 | CRPS | 99.3 | 98.5 | 95.7 | 99.1 | 94.5 | 96.8 | 90.2 | 87.5 | 85.7 | 89.3 | 78.6 | 70.4 |
| P18 | CRPS | 89.1 | 96.2 | 91.5 | 95.7 | 74.5 | 88.7 | 75.4 | 80.4 | 57.1 | 78.6 | 60.7 | 81.5 |
| P19 | CRPS | 97.8 | 100.0 | 99.3 | 97.4 | 90.9 | 96.8 | 90.2 | 98.2 | 64.3 | 96.4 | 64.3 | 85.2 |
| P20 | CRPS | 98.5 | 100.0 | 98.6 | 99.1 | 87.3 | 87.1 | 86.9 | 91.1 | 75.0 | 78.6 | 71.4 | 77.8 |
| P21 | CRPS | 99.3 | 98.5 | 97.9 | 100.0 | 85.5 | 95.2 | 86.9 | 92.9 | 53.6 | 89.3 | 67.9 | 85.2 |
| P22 | CRPS | 99.3 | 96.2 | 99.3 | 100.0 | 78.2 | 100.0 | 98.4 | 100.0 | 78.6 | 85.7 | 82.1 | 92.6 |
| Mean (group) |  | 95.8 | 97.4 | 96.3 | 97.8 | 83.7 | 92.0 | 88.0 | 91.1 | 71.9 | 83.6 | 75.6 | 81.9 |
| SD (group) |  | 4.7 | 3.1 | 2.8 | 2.6 | 6.8 | 7.2 | 7.0 | 6.3 | 10.6 | 11.9 | 10.0 | 10.0 |
| **P23** | HC | 100.0 | 100.0 | 99.3 | 100.0 | 85.5 | 91.9 | 91.8 | 91.1 | 67.9 | 75.0 | 64.3 | 74.1 |
| **P24** | HC | 92.7 | 94.7 | 94.3 | 94.8 | 80.0 | 82.3 | 86.9 | 87.5 | 64.3 | 71.4 | 78.6 | 70.4 |
| **P25** | HC | 96.4 | 95.5 | 92.9 | 96.6 | 83.6 | 87.1 | 90.2 | 94.6 | 71.4 | 67.9 | 64.3 | 74.1 |
| **P26** | HC | 100.0 | 98.3 | 92.7 | 95.5 | 82.0 | 83.9 | 90.9 | 90.3 | 78.6 | 96.3 | 67.9 | 57.1 |
| **P27** | HC | 94.9 | 95.5 | 91.5 | 99.1 | 76.4 | 87.1 | 86.9 | 85.7 | 78.6 | 89.3 | 78.6 | 63.0 |
| **P28** | HC | 98.5 | 97.7 | 92.9 | 98.3 | 80.0 | 79.0 | 93.4 | 87.5 | 82.1 | 75.0 | 75.0 | 88.9 |
| **P29** | HC | 99.3 | 99.1 | 100.0 | 100.0 | 100.0 | 98.2 | 100.0 | 96.8 | 89.3 | 92.6 | 89.3 | 96.4 |
| **P30** | HC | 100.0 | 100.0 | 99.3 | 100.0 | 85.5 | 93.5 | 91.8 | 83.9 | 82.1 | 82.1 | 89.3 | 88.9 |
| **P31** | HC | 98.5 | 100.0 | 97.9 | 97.4 | 81.8 | 98.4 | 83.6 | 87.5 | 67.9 | 82.1 | 67.9 | 81.5 |
| **P32** | HC | 100.0 | 100.0 | 100.0 | 99.2 | 95.1 | 92.9 | 94.5 | 96.8 | 89.3 | 100.0 | 71.4 | 92.9 |
| **P33** | HC | 99.3 | 100.0 | 98.6 | 100.0 | 89.1 | 90.3 | 88.5 | 92.9 | 85.7 | 71.4 | 89.3 | 81.5 |
| **P34** | HC | 95.6 | 93.9 | 87.9 | 91.4 | 72.7 | 83.9 | 75.4 | 91.1 | 67.9 | 60.7 | 75.0 | 70.4 |
| **P35** | HC | 98.5 | 100.0 | 97.9 | 100.0 | 92.7 | 96.8 | 85.2 | 100.0 | 85.7 | 100.0 | 89.3 | 96.3 |
| **P36** | HC | 94.9 | 93.9 | 92.9 | 97.4 | 74.5 | 85.5 | 78.7 | 85.7 | 57.1 | 82.1 | 60.7 | 88.9 |
| **P37** | HC | 98.6 | 99.1 | 97.8 | 98.5 | 96.7 | 92.9 | 92.7 | 87.1 | 85.7 | 92.6 | 85.7 | 96.4 |
| **P38** | HC | 91.2 | 93.9 | 87.9 | 90.5 | 80.0 | 85.5 | 77.0 | 92.9 | 92.9 | 78.6 | 89.3 | 70.4 |
| **P39** | HC | 98.5 | 100.0 | 100.0 | 100.0 | 90.9 | 100.0 | 95.1 | 100.0 | 85.7 | 100.0 | 96.4 | 100.0 |
| **P40** | HC | 100.0 | 100.0 | 99.3 | 99.1 | 92.7 | 96.8 | 96.7 | 100.0 | 85.7 | 92.9 | 89.3 | 92.6 |
| **P41** | HC | 98.5 | 98.5 | 97.9 | 100.0 | 80.0 | 87.1 | 80.3 | 87.5 | 78.6 | 85.7 | 82.1 | 81.5 |
| **P42** | HC | 96.5 | 99.1 | 100.0 | 100.0 | 98.4 | 100.0 | 94.5 | 98.4 | 92.9 | 96.3 | 78.6 | 96.4 |
| **P43** | HC | 94.9 | 98.5 | 97.2 | 94.0 | 81.8 | 88.7 | 88.5 | 85.7 | 75.0 | 96.4 | 71.4 | 81.5 |
| **P44** | HC | 94.2 | 98.5 | 97.9 | 99.1 | 83.6 | 91.9 | 100.0 | 98.2 | 89.3 | 100.0 | 78.6 | 100.0 |
| **Mean**<br>**(group)** |  | 97.3 | 98.0 | 96.2 | 97.8 | 85.6 | 90.6 | 89.2 | 91.9 | 79.7 | 85.8 | 78.7 | 83.8 |
| **SD (group)** |  | 2.6 | 2.3 | 3.9 | 2.9 | 7.8 | 6.1 | 7.0 | 5.4 | 9.9 | 11.9 | 10.1 | 12.5 |
| **Mann-Whitney U test** |  |  |  |  |  |  |  |  |  |  |  |  |  |
| **Effect size: Z statistic** |  | -0.82 | -0.63 | 0.72 | 0.23 | -0.72 | 0.79 | -0.72 | -0.24 | -2.36 | -0.77 | -1.08 | -0.95 |

|  |  |  |  |  |  |  |  |  |  |  |  |  |
| --- | --- | --- | --- | --- | --- | --- | --- | --- | --- | --- | --- | --- |
| **P value (asymptotic 2-tailed)** | 0.412 | 0.531 | 0.472 | 0.815 | 0.474 | 0.432 | 0.474 | 0.811 | 0.019 | 0.438 | 0.282 | 0.345 |
| **Corrected p value** | 4.939 | 6.377 | 5.669 | 9.781 | 5.683 | 5.187 | 5.686 | 9.733 | 0.222 | 5.260 | 3.383 | 4.136 |

**Supplementary Table 5:**
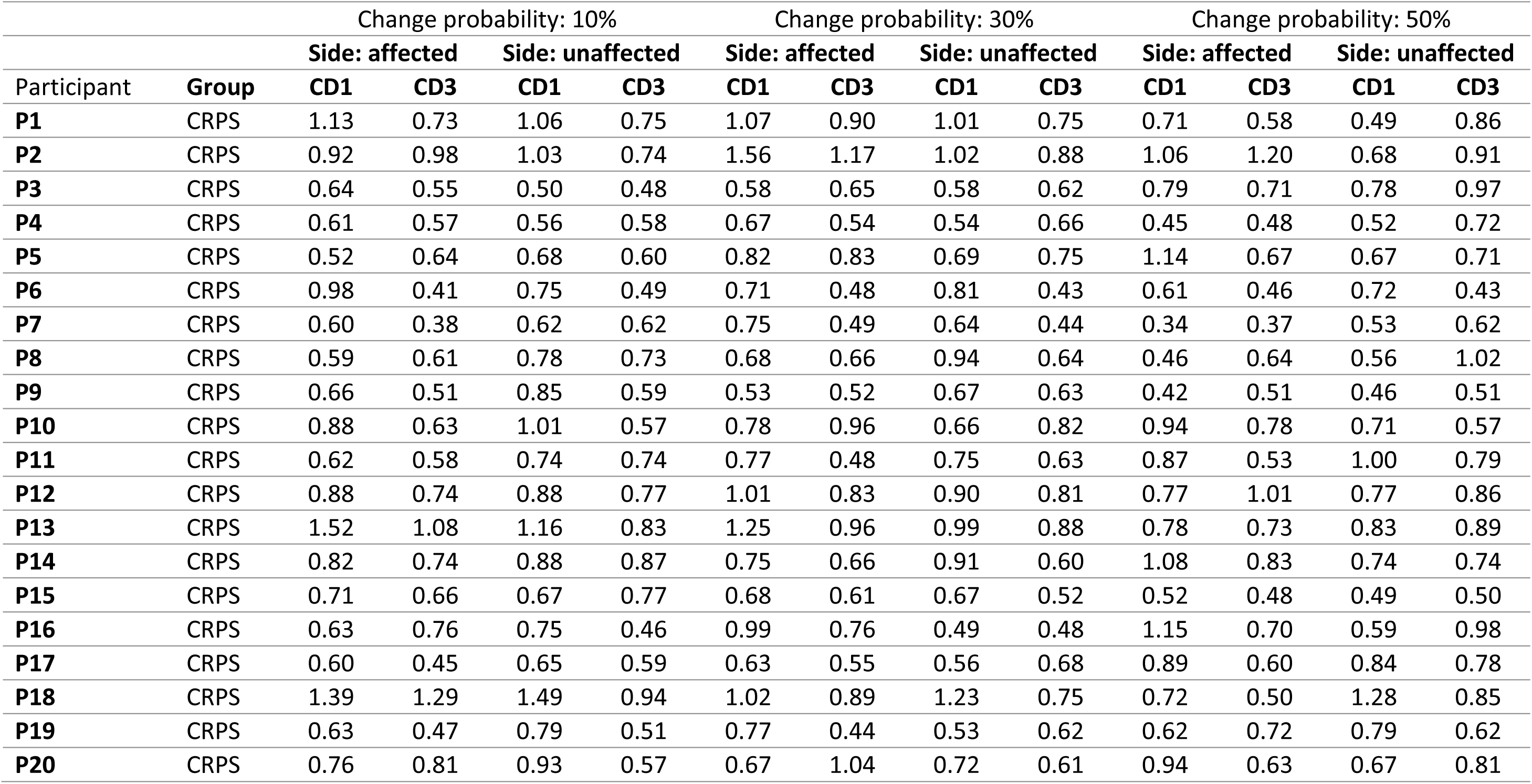
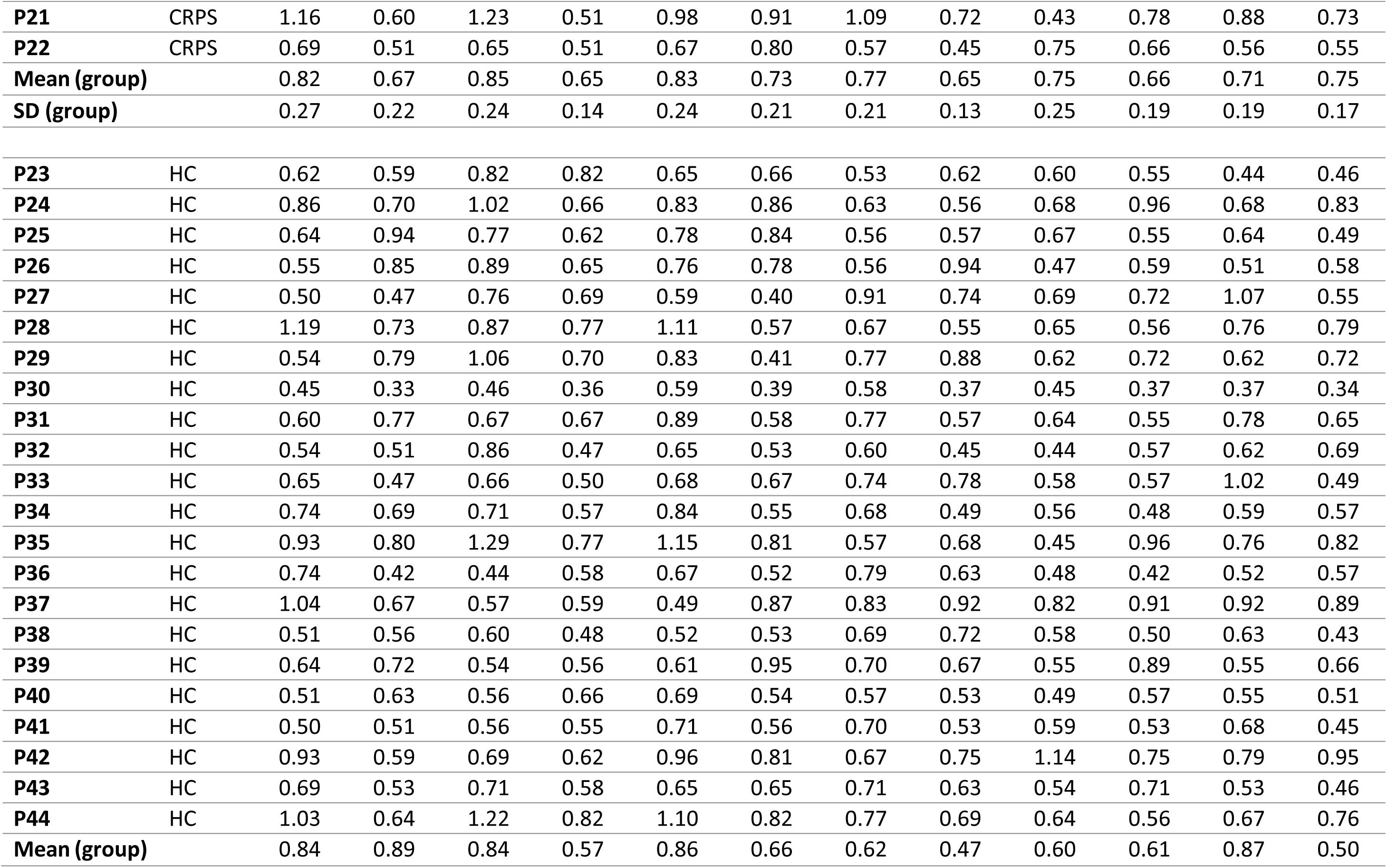
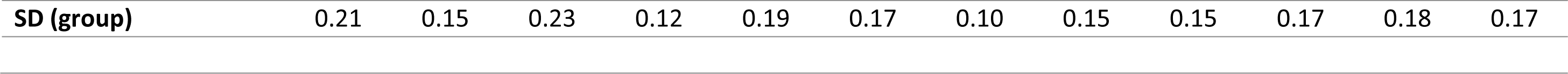
Tactile Spatial Oddball Task (TSOT) response time data: individual means.

| Participant | Group | Change probability: 10% |  |  |  | Change probability: 30% |  |  |  | Change probability: 50% |  |  |  |
| --- | --- | --- | --- | --- | --- | --- | --- | --- | --- | --- | --- | --- | --- |
|  |  | Side: affected |  | Side: unaffected |  | Side: affected |  | Side: unaffected |  | Side: affected |  | Side: unaffected |  |
|  |  | CD1 | CD3 | CD1 | CD3 | CD1 | CD3 | CD1 | CD3 | CD1 | CD3 | CD1 | CD3 |
| **P1** | CRPS | 1.13 | 0.73 | 1.06 | 0.75 | 1.07 | 0.90 | 1.01 | 0.75 | 0.71 | 0.58 | 0.49 | 0.86 |
| **P2** | CRPS | 0.92 | 0.98 | 1.03 | 0.74 | 1.56 | 1.17 | 1.02 | 0.88 | 1.06 | 1.20 | 0.68 | 0.91 |
| **P3** | CRPS | 0.64 | 0.55 | 0.50 | 0.48 | 0.58 | 0.65 | 0.58 | 0.62 | 0.79 | 0.71 | 0.78 | 0.97 |
| **P4** | CRPS | 0.61 | 0.57 | 0.56 | 0.58 | 0.67 | 0.54 | 0.54 | 0.66 | 0.45 | 0.48 | 0.52 | 0.72 |
| **P5** | CRPS | 0.52 | 0.64 | 0.68 | 0.60 | 0.82 | 0.83 | 0.69 | 0.75 | 1.14 | 0.67 | 0.67 | 0.71 |
| **P6** | CRPS | 0.98 | 0.41 | 0.75 | 0.49 | 0.71 | 0.48 | 0.81 | 0.43 | 0.61 | 0.46 | 0.72 | 0.43 |
| **P7** | CRPS | 0.60 | 0.38 | 0.62 | 0.62 | 0.75 | 0.49 | 0.64 | 0.44 | 0.34 | 0.37 | 0.53 | 0.62 |
| **P8** | CRPS | 0.59 | 0.61 | 0.78 | 0.73 | 0.68 | 0.66 | 0.94 | 0.64 | 0.46 | 0.64 | 0.56 | 1.02 |
| **P9** | CRPS | 0.66 | 0.51 | 0.85 | 0.59 | 0.53 | 0.52 | 0.67 | 0.63 | 0.42 | 0.51 | 0.46 | 0.51 |
| **P10** | CRPS | 0.88 | 0.63 | 1.01 | 0.57 | 0.78 | 0.96 | 0.66 | 0.82 | 0.94 | 0.78 | 0.71 | 0.57 |
| **P11** | CRPS | 0.62 | 0.58 | 0.74 | 0.74 | 0.77 | 0.48 | 0.75 | 0.63 | 0.87 | 0.53 | 1.00 | 0.79 |
| **P12** | CRPS | 0.88 | 0.74 | 0.88 | 0.77 | 1.01 | 0.83 | 0.90 | 0.81 | 0.77 | 1.01 | 0.77 | 0.86 |
| **P13** | CRPS | 1.52 | 1.08 | 1.16 | 0.83 | 1.25 | 0.96 | 0.99 | 0.88 | 0.78 | 0.73 | 0.83 | 0.89 |
| **P14** | CRPS | 0.82 | 0.74 | 0.88 | 0.87 | 0.75 | 0.66 | 0.91 | 0.60 | 1.08 | 0.83 | 0.74 | 0.74 |
| **P15** | CRPS | 0.71 | 0.66 | 0.67 | 0.77 | 0.68 | 0.61 | 0.67 | 0.52 | 0.52 | 0.48 | 0.49 | 0.50 |
| **P16** | CRPS | 0.63 | 0.76 | 0.75 | 0.46 | 0.99 | 0.76 | 0.49 | 0.48 | 1.15 | 0.70 | 0.59 | 0.98 |
| **P17** | CRPS | 0.60 | 0.45 | 0.65 | 0.59 | 0.63 | 0.55 | 0.56 | 0.68 | 0.89 | 0.60 | 0.84 | 0.78 |
| **P18** | CRPS | 1.39 | 1.29 | 1.49 | 0.94 | 1.02 | 0.89 | 1.23 | 0.75 | 0.72 | 0.50 | 1.28 | 0.85 |
| **P19** | CRPS | 0.63 | 0.47 | 0.79 | 0.51 | 0.77 | 0.44 | 0.53 | 0.62 | 0.62 | 0.72 | 0.79 | 0.62 |
| **P20** | CRPS | 0.76 | 0.81 | 0.93 | 0.57 | 0.67 | 1.04 | 0.72 | 0.61 | 0.94 | 0.63 | 0.67 | 0.81 |
| **P21** | CRPS | 1.16 | 0.60 | 1.23 | 0.51 | 0.98 | 0.91 | 1.09 | 0.72 | 0.43 | 0.78 | 0.88 | 0.73 |
| **P22** | CRPS | 0.69 | 0.51 | 0.65 | 0.51 | 0.67 | 0.80 | 0.57 | 0.45 | 0.75 | 0.66 | 0.56 | 0.55 |
| **Mean (group)** |  | 0.82 | 0.67 | 0.85 | 0.65 | 0.83 | 0.73 | 0.77 | 0.65 | 0.75 | 0.66 | 0.71 | 0.75 |
| **SD (group)** |  | 0.27 | 0.22 | 0.24 | 0.14 | 0.24 | 0.21 | 0.21 | 0.13 | 0.25 | 0.19 | 0.19 | 0.17 |
| **P23** | HC | 0.62 | 0.59 | 0.82 | 0.82 | 0.65 | 0.66 | 0.53 | 0.62 | 0.60 | 0.55 | 0.44 | 0.46 |
| **P24** | HC | 0.86 | 0.70 | 1.02 | 0.66 | 0.83 | 0.86 | 0.63 | 0.56 | 0.68 | 0.96 | 0.68 | 0.83 |
| **P25** | HC | 0.64 | 0.94 | 0.77 | 0.62 | 0.78 | 0.84 | 0.56 | 0.57 | 0.67 | 0.55 | 0.64 | 0.49 |
| **P26** | HC | 0.55 | 0.85 | 0.89 | 0.65 | 0.76 | 0.78 | 0.56 | 0.94 | 0.47 | 0.59 | 0.51 | 0.58 |
| **P27** | HC | 0.50 | 0.47 | 0.76 | 0.69 | 0.59 | 0.40 | 0.91 | 0.74 | 0.69 | 0.72 | 1.07 | 0.55 |
| **P28** | HC | 1.19 | 0.73 | 0.87 | 0.77 | 1.11 | 0.57 | 0.67 | 0.55 | 0.65 | 0.56 | 0.76 | 0.79 |
| **P29** | HC | 0.54 | 0.79 | 1.06 | 0.70 | 0.83 | 0.41 | 0.77 | 0.88 | 0.62 | 0.72 | 0.62 | 0.72 |
| **P30** | HC | 0.45 | 0.33 | 0.46 | 0.36 | 0.59 | 0.39 | 0.58 | 0.37 | 0.45 | 0.37 | 0.37 | 0.34 |
| **P31** | HC | 0.60 | 0.77 | 0.67 | 0.67 | 0.89 | 0.58 | 0.77 | 0.57 | 0.64 | 0.55 | 0.78 | 0.65 |
| **P32** | HC | 0.54 | 0.51 | 0.86 | 0.47 | 0.65 | 0.53 | 0.60 | 0.45 | 0.44 | 0.57 | 0.62 | 0.69 |
| **P33** | HC | 0.65 | 0.47 | 0.66 | 0.50 | 0.68 | 0.67 | 0.74 | 0.78 | 0.58 | 0.57 | 1.02 | 0.49 |
| **P34** | HC | 0.74 | 0.69 | 0.71 | 0.57 | 0.84 | 0.55 | 0.68 | 0.49 | 0.56 | 0.48 | 0.59 | 0.57 |
| **P35** | HC | 0.93 | 0.80 | 1.29 | 0.77 | 1.15 | 0.81 | 0.57 | 0.68 | 0.45 | 0.96 | 0.76 | 0.82 |
| **P36** | HC | 0.74 | 0.42 | 0.44 | 0.58 | 0.67 | 0.52 | 0.79 | 0.63 | 0.48 | 0.42 | 0.52 | 0.57 |
| **P37** | HC | 1.04 | 0.67 | 0.57 | 0.59 | 0.49 | 0.87 | 0.83 | 0.92 | 0.82 | 0.91 | 0.92 | 0.89 |
| **P38** | HC | 0.51 | 0.56 | 0.60 | 0.48 | 0.52 | 0.53 | 0.69 | 0.72 | 0.58 | 0.50 | 0.63 | 0.43 |
| **P39** | HC | 0.64 | 0.72 | 0.54 | 0.56 | 0.61 | 0.95 | 0.70 | 0.67 | 0.55 | 0.89 | 0.55 | 0.66 |
| **P40** | HC | 0.51 | 0.63 | 0.56 | 0.66 | 0.69 | 0.54 | 0.57 | 0.53 | 0.49 | 0.57 | 0.55 | 0.51 |
| **P41** | HC | 0.50 | 0.51 | 0.56 | 0.55 | 0.71 | 0.56 | 0.70 | 0.53 | 0.59 | 0.53 | 0.68 | 0.45 |
| **P42** | HC | 0.93 | 0.59 | 0.69 | 0.62 | 0.96 | 0.81 | 0.67 | 0.75 | 1.14 | 0.75 | 0.79 | 0.95 |
| **P43** | HC | 0.69 | 0.53 | 0.71 | 0.58 | 0.65 | 0.65 | 0.71 | 0.63 | 0.54 | 0.71 | 0.53 | 0.46 |
| **P44** | HC | 1.03 | 0.64 | 1.22 | 0.82 | 1.10 | 0.82 | 0.77 | 0.69 | 0.64 | 0.56 | 0.67 | 0.76 |
| **Mean (group)** |  | 0.84 | 0.89 | 0.84 | 0.57 | 0.86 | 0.66 | 0.62 | 0.47 | 0.60 | 0.61 | 0.87 | 0.50 |

|  |  |  |  |  |  |  |  |  |  |  |  |  |
| --- | --- | --- | --- | --- | --- | --- | --- | --- | --- | --- | --- | --- |
| **SD (group)** | 0.21 | 0.15 | 0.23 | 0.12 | 0.19 | 0.17 | 0.10 | 0.15 | 0.15 | 0.17 | 0.18 | 0.17 |

**Supplementary Table 6:**
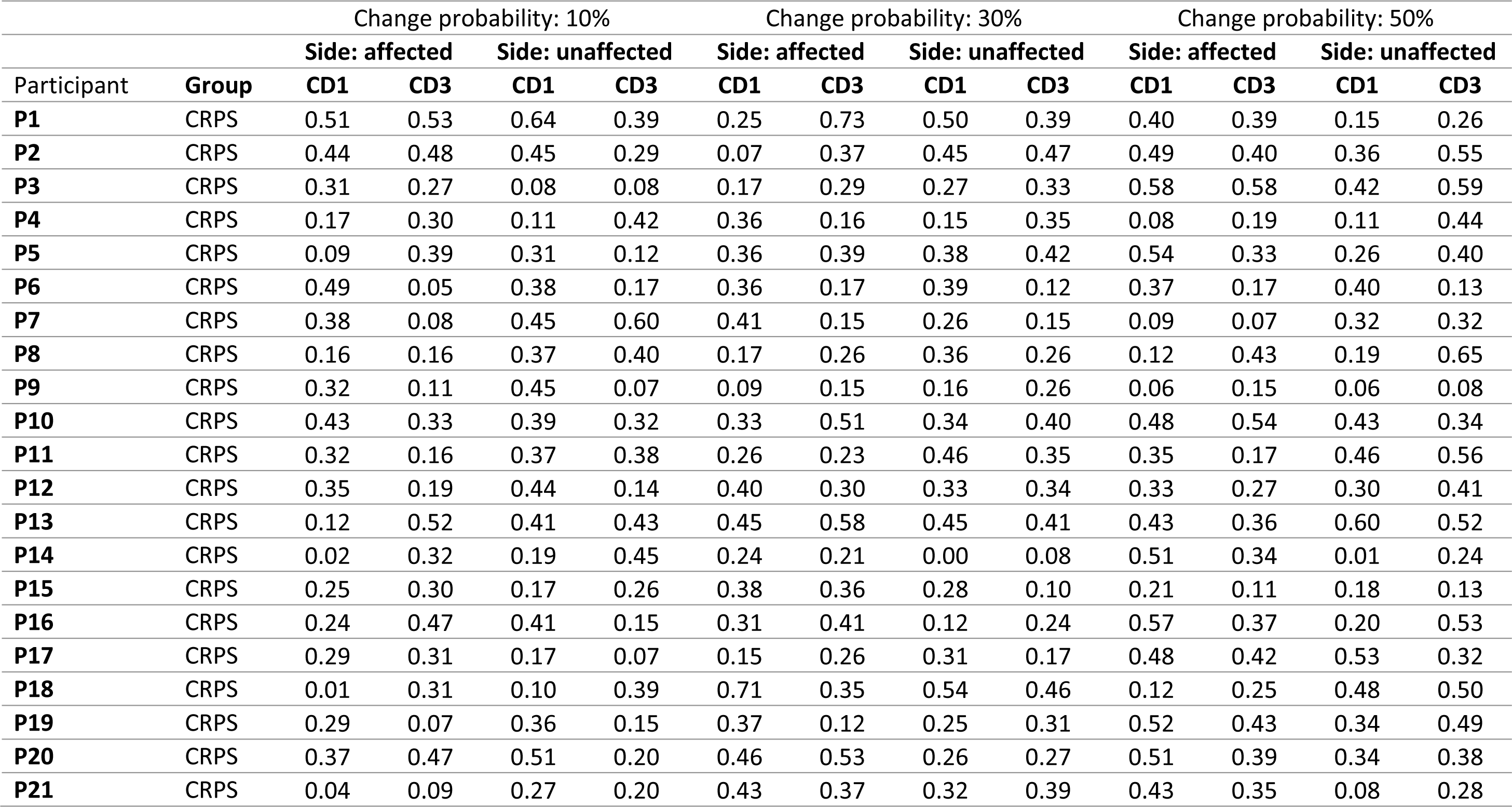
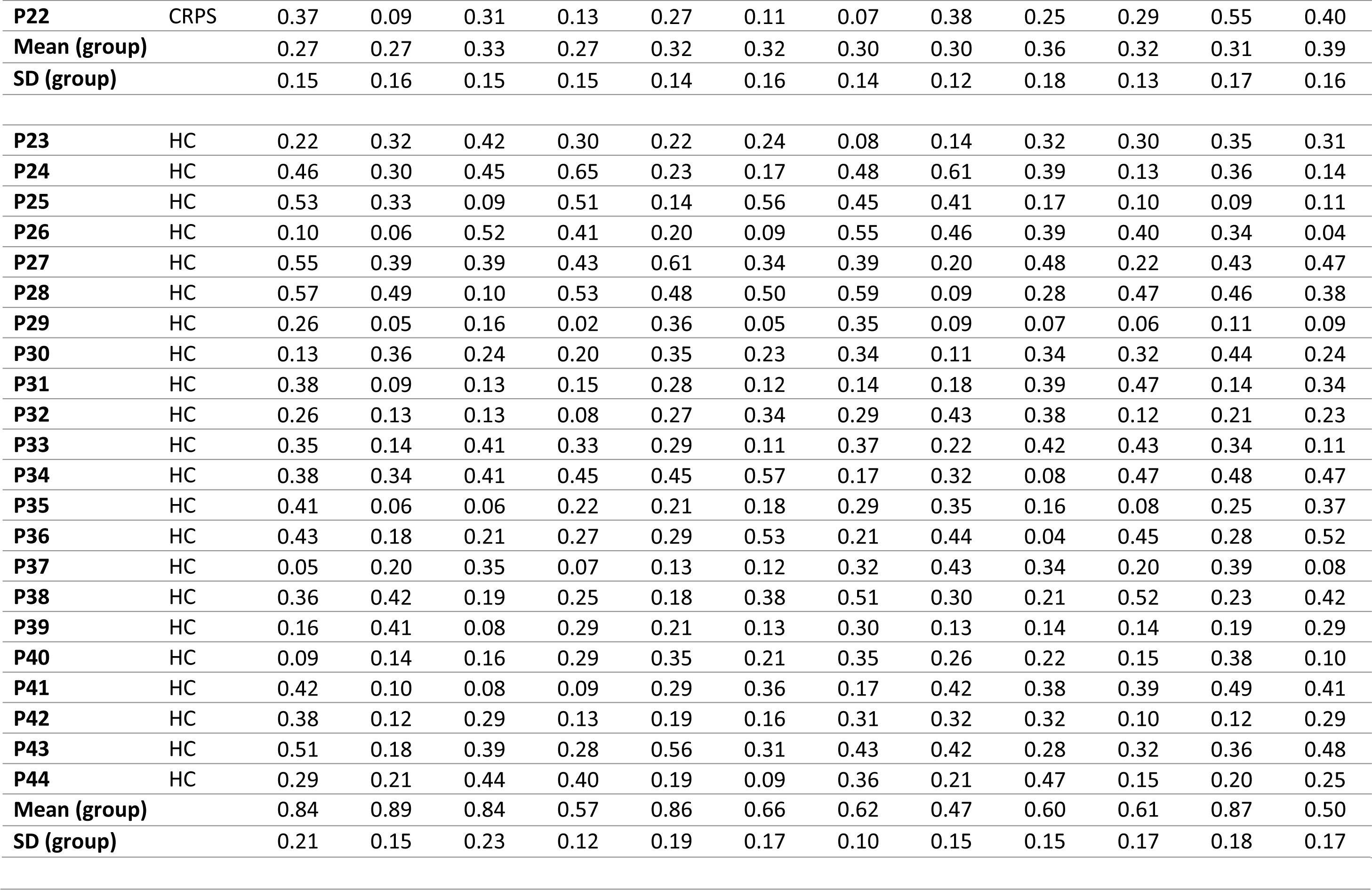
Tactile Spatial Oddball Task (TSOT) response time data: individual standard deviations.

**Supplementary Table 7:** Mixed ANOVA on log(RT) data from Tactile Spatial Oddball Task (TSOT)

|  | Effect | n | df1 | df2 | F | p | Partial<br>Eta<br>Squared |
| --- | --- | --- | --- | --- | --- | --- | --- |
| **ANOVA on**<br>**means**<br>**(over trials**<br>**per**<br>**condition /**<br>**individual)** | CP † | 44 | 1.54 | 64.73 | 3.54 | **0.046** | 0.08 |
|  | Side | 44 | 1 | 42 | 0.00 | 0.972 | 0.00 |
|  | CD | 44 | 1 | 42 | 30.08 | **0.000** | 0.42 |
|  | CP * Side † | 44 | 2 | 84 | 5.06 | **0.008** | 0.11 |
|  | CP * CD † | 44 | 2 | 84 | 8.18 | **0.001** | 0.16 |
|  | Side * CD | 44 | 1 | 42 | 0.02 | 0.894 | 0.00 |
|  | CP * Side * CD † | 44 | 2 | 84 | 1.33 | 0.271 | 0.03 |
|  | CP * Group † | 30 | 1.54 | 64.73 | 0.05 | 0.919 | 0.00 |
|  | Side * Group | 30 | 1 | 28 | 0.01 | 0.910 | 0.00 |
|  | CD * Group | 30 | 1 | 28 | 1.73 | 0.199 | 0.06 |
|  | CP * Side * Group † | 30 | 2 | 56 | 0.58 | 0.555 | 0.02 |
|  | CP * CD * Group † | 30 | 2 | 56 | 0.30 | 0.719 | 0.01 |
|  | Side * CD * Group | 30 | 1 | 28 | 0.60 | 0.443 | 0.02 |
|  | CP * Side * CD * Group † | 30 | 2 | 56 | 1.11 | 0.333 | 0.04 |
|  | Group | 30 | 1 | 28 | 0.51 | 0.479 | 0.02 |
| **ANOVA on**<br>**standard**<br>**deviations** | CP | 44 | 1.54 | 64.73 | 2.01 | 0.140 | 0.05 |
|  | Side | 44 | 1 | 42 | 0.73 | 0.399 | 0.02 |
|  | CD | 44 | 1 | 42 | 1.60 | 0.213 | 0.04 |
| (over trials<br>per<br>condition /<br>individual) | CP * Side | 44 | 2 | 84 | 0.00 | 0.996 | 0.00 |
|  | CP * CD | 44 | 2 | 84 | 0.80 | 0.452 | 0.02 |
|  | Side * CD | 44 | 1 | 42 | 1.72 | 0.197 | 0.04 |
|  | CP * Side * CD | 44 | 2 | 84 | 0.83 | 0.440 | 0.02 |
|  | CP * Group | 30 | 1.54 | 64.74 | 1.97 | 0.150 | 0.07 |
|  | Side * Group | 30 | 1 | 28 | 0.21 | 0.651 | 0.01 |
|  | CD * Group | 30 | 1 | 28 | 0.19 | 0.670 | 0.01 |
|  | CP * Side * Group | 30 | 2 | 56 | 1.39 | 0.258 | 0.05 |
|  | CP * CD * Group | 30 | 2 | 56 | 0.07 | 0.934 | 0.00 |
|  | Side * CD * Group | 30 | 1 | 28 | 0.06 | 0.813 | 0.00 |
|  | CP * Side * CD * Group | 30 | 2 | 56 | 6.17 | **0.004** | 0.18 |
|  | Group | 30 | 1 | 28 | 0.05 | 0.832 | 0.02 |

CP: Change Probability
CD: Change Distance
† Greenhouse-Geisser-corrected due to failing Mauchley's test for sphericity

### Computational modelling rationale

Computational modelling of TSOT response times employed the Hierarchical Gaussian Filter (HGF) toolbox v5.0 with additional customised model functions. The HGF (Mathys et al., 2014) is a mathematical model of a particular type of predictive coding that implements hierarchical Bayesian inference using a variational Bayes approximation scheme, consistent with the “free-energy” formulation of predictive coding (Friston and Kiebel, 2009). The HGF is particularly suitable for modelling the effects of volatile (non-stationary) environments on behaviour, in that it acts as a generative model of the environment – i.e. a model of how the environment is structured to cause experienced sensations. Such generative models can operate as a “forward model” to simulate behaviour (in response to pre-defined sensory inputs), or as an “inverse model” to both infer the nature of sensations and learn the parameters of the model simultaneously (in response to pre-defined sensory inputs and observed participant responses). More specifically for our experiment, we employed the HGF as a model of (1) how the brain might represent the environmental contingencies that could give rise to changes in tactile stimulus locations (the “generative” aspect of the model), (2) how the brain might use those representations to make inferences about whether a location change has taken place (the “inferential” aspect), and (3) how the brain might update its parameters in light of new sensory information (the “learning” aspect). The model combines inference and learning via minimisation of prediction error (mismatches between predicted and actual sensory inputs). In sum, this provides a model of how the brain might perform approximate Bayesian inference to infer spatial changes in tactile sensations.

While many Bayesian models of perception are “ideal observer” models and assume optimality in how sensory information updated hidden representations of the environment (Maloney and Zhang, 2010), the HGF instead assumes that the optimality of an update may vary between participants due to differences in prior beliefs about the higher-order structure of the environment (e.g. change probabilities and how they evolve over time) (Mathys et al., 2014). These prior beliefs are partly a product of two factors: prior beliefs placed on the models as experimenters (i.e. the model “priors” in a Bayesian sense), and the history of sensory information that updates those priors during the experiment. Importantly, there is a third factor that the model assumes can vary over participants: parameters of the model that are fixed over the whole experiment (over all sensory inputs) and that modulate the “gain” of representation updates. These parameters account for inter-individual differences in learning from sensory inputs, and are estimated by fitting the model to actual participant behaviour to infer those parameters, using a variational Bayesian optimisation algorithm. Ultimately, such a model is capable of describing behaviour that is subjectively optimal (in relation to the participant’s prior beliefs) but might be objectively maladaptive (resulting in inaccurate or delayed inferences, relative to an ideal observer) (Mathys et al., 2014).

Theoretically, the model parameters that determine learning may relate to specific physiological processes, such as the neuromodulation of synaptic plasticity (Bogacz, 2017). The HGF uses closed-form update equations to calculate the posterior expectations of all hidden states (representations) in the model. These are efficient computations that allow for real-time learning and have been shown to be biologically plausible (Bogacz, 2017). Furthermore, the form of these update equations is similar to those of Rescorla–Wagner (RW) learning (Rescorla and Wagner, 1972), a widely successful framework for associative learning, in that inference proceeds by updating representations according to prediction errors. The HGF departs from RW learning in that prediction error updates are precision-weighted, i.e. weighted according to the relative confidence/certainty in lower-level information (e.g. sensory inputs) over higher-level representations (e.g. predictions of sensory inputs) (Mathys et al., 2014). In this regard, the HGF accounts for different types of uncertainty, namely sensory uncertainty (uncertainty in sensory inputs, a.k.a. irreducible uncertainty), expected uncertainty (uncertainty in learnt predictions of sensory inputs), and environmental volatility (uncertainty in how predictions should change over time). These three levels of uncertainty correspond to the 3 levels of the HGF model (Fig. 2b of main paper).

In addition to a perceptual model of the environment represented by these three levels of the HGF, in order to fit the parameters of the model as experimenters, we also need a response model that maps representations within the model onto observed behaviour (Mathys et al., 2014). In the absence of a response model, we can still invert the model (estimate the parameters) from sensory inputs alone (i.e. the experimental design), but this would only provide us with state representations and parameters that represent an ideal observer (an observer who computes the least “surprise” about sensory inputs). To estimate subject-specific parameters (as indices of each participant’s deviation from optimality) requires the introduction of a response model. Just as for the perceptual model, the response model also contains subject-specific response parameters (Fig. 2b of main paper).

### HGF implementation details

Our implementation of the HGF differs from previously published treatments (Mathys et al., 2014) in the following ways. Firstly, with regard to the perceptual model, we tested 3 perceptual models that provided variation on the common model structure shown in Fig. 2b (main paper). The 3 models varied according to the optional addition of two further parameters at the sensory input level (λ_1_, λ_2_) that were designed to better capture variability in behaviour resulting from condition effects on sensory uncertainty. Each additional parameter acted as a multiplier on the α_0_ parameter, in order to capture differences in estimated sensory uncertainty arising from two factors in the design: firstly, λ_1_ captured within-subject variability in sensory uncertainty arising from the digit change distance (CD1 vs. CD3), and secondly, λ_2_ acted to capture additional variability arising from the hand stimulated (affected vs. unaffected). The 3 perceptual models consisted of no additional parameters (p-model 1), an additional λ_1_ parameter (p-model 2), or the addition of both λ_1_ and λ_2_ (p-model 3). In other words, to take the example of the most complex perceptual model (p-model 3), inverting the model allowed estimation, for each subject, the overall sensory uncertainty (α_0_), how α_0_ varies by digit change distance (λ_1_) and how α_0_ varied by hand stimulated (λ_2_).

Mathematically, these were implemented as follows for perceptual model 3. Here we consider the variance that is common to all conditions as, as well as multipliers from two orthogonal factors in the design, a “Side” condition (*λ_side_*) and a “Change Distance (CD)” condition (*λ_dc_*). Regarding the likelihood function within the HGF model, now *u* is conditionally dependent not just on state *x*, but also on conditions 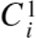(CD) and 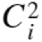(side) (where *i* ∈ 007B;0,1| represents two levels of each condition):

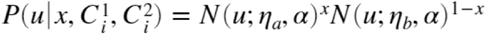

where

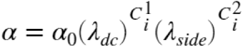

Orthogonally to the perceptual models, we tested a total of 6 response models, all of which sought to link states common to all perceptual models to log-RT data. Each response model is a simple linear model, each with either 3 or 4 parameters (β_1_, β_2_, β_3_, plus or minus β_4_) that provide weights to a different set of states in the perceptual model, plus a further parameter describing a constant component (β_0_). The (Gaussian) error term in this linear model is denoted by ζ, which acts as an inverse decision temperature (greater error = behaviour more stochastic / less determined by the model). The variables within each of the first four response models were different types of PE, based on the hypothesis that RTs would be slower to unexpected spatial changes:
r-model 1: Signed, unweighted, PEs at each of the 3 p-model levels (δ_1_, δ_2_, δ_3_)

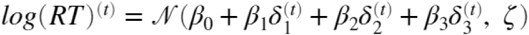

r-model 2: Absolute, unweighted, PEs at each of the 3 p-model levels (|δ_1_|, |δ_2_|, |δ_3_|)

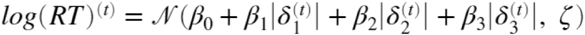

r-model 3: Signed, precision-weighted, PEs at each of the 3 p-model levels (ε_1_, ε_2_, ε_3_)

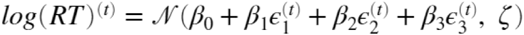

r-model 4: Absolute, precision-weighted, PEs at each of the 3 p-model levels (|ε_1_|, |ε_2_|, |ε_3_|)

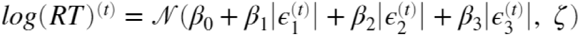

The final two response models consisted of variables related to uncertainty at each level of the perceptual model, for consistency with previous literature showing that these variables are linked to RT on perceptual tasks (Lawson et al., 2017; Marshall et al., 2016). These models provided a benchmark against which to test whether the above response models consisting of PE terms provide better or worse performance in predicting log-RTs.

r-model 5: Uncertainty (state variance) at each of the 3 p-model levels model (*unc*_1_, *unc*_2_ and *unc*_3_). We used versions of uncertainty published previously (Lawson et al., 2017; Marshall et al., 2016): 1^st^ level Bernoulli variance (σ_1_), 2^nd^ level inferential variance (sigmoid transformation of σ_2_), and 3^rd^ level phasic volatility (sigmoid transformation of the exponent of µ_3_).

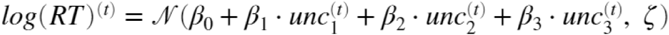

r-model 6: Uncertainty (state variance) at each of the 3 p-model levels as outlined for r-model 5, but with the addition of a fourth variable and associated parameter known as “information surprise” at level 1 (described in (Lawson et al., 2017; Marshall et al., 2016).

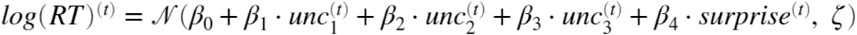

where independent variables are defined as:

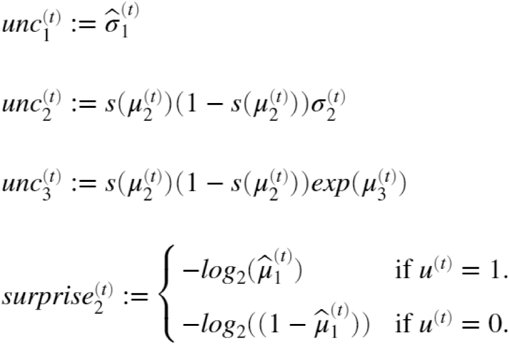

All 18 models (3 perceptual models x 6 response models) were estimated using the same priors, except where differences in model structure did not require certain priors. Firstly, priors were required for initial states prior to the first trial, which were decided a priori and fixed (see Supplementary Table 8). Secondly, subject-specific parameters were either decided a priori (in the case of λ_1_ and λ_2_, and response model parameters) or by using the parameter estimates from optimising a Bayes-optimal model (for α_0_, ω and θ).

**Supplementary Table 8:** HGF model priors.

| Parameter type | Parameter | Prior mean | Prior variance (0 = fixed) |
| --- | --- | --- | --- |
| Perceptual model initial states | $\eta_0$ | 0 | 0 |
| | $\eta_1$ | 1 | 0 |
| | $\mu_0(x_1)$ | none | none |
| | $\sigma_0(x_1)$ | none | none |
| | $\mu_0(x_2)$ | 0 | 0 |
| | $\sigma_0(x_2)$ | 0.5 | 0 |
| | $\mu_0(x_3)$ | 1 | 0 |
| | $\sigma_0(x_3)$ | 1 | 0 |
| | $\rho(x_1)$ | none | none |
| | $\rho(x_2)$ | 0 | 0 |
| | $\rho(x_3)$ | 0 | 0 |
| | $\kappa(x_1)$ | none | none |
| | $\kappa(x_2)$ | 1 | 0 |
| Perceptual model subject-specific | $\alpha_0$ | 0.0925 | 1 |
| | $\lambda_1$ | 1 | 1† |
| | $\lambda_2$ | 1 | 1†† |
| | $\omega$ | -5.1826 | 16 |
| | $\theta$ | -6.8176 | 16 |
| Response model subject-specific | $\beta_0$ | 0.5 ††† | 4 |
| | $\beta_{1-4}$ | 0 | 4 |
| | $\zeta$ | log(20) | 2 |

† Parameter variances were set to zero for perceptual models 1 and 2. †† Parameter variances were set to zero for perceptual model 2. ††† Expected mean response time

**Supplementary Table 9:** HGF model posterior estimates by group.

| Estimate type | Estimate | CRPS group (n=15) |  | HC group (n=15) |  | Group effects |  |
| --- | --- | --- | --- | --- | --- | --- | --- |
|  |  | median | IQR | median | IQR | Z | p |
| Perceptual model subject-specific parameter | $\alpha_0$ | 0.01 | 0.08 | -0.04 | 0.11 | 1.29 | 0.199 |
| | $\lambda_1$ | -0.09 | 0.38 | -0.04 | 0.46 | -0.25 | 0.803 |
| | $\lambda_2$ | -0.08 | 0.23 | -0.04 | 0.32 | -0.25 | 0.803 |
| | $\omega$ | 0.79 | 1.37 | -0.41 | 1.31 | 2.99 | **0.003** |
| | $\theta$ | -1.04 | 1.23 | 1.52 | 1.33 | -3.98 | **<0.001** |
| | $\beta_0$ | 0.00 | 0.27 | 0.01 | 0.16 | 0.71 | 0.481 |
| Response model<br>subject-specific<br>parameter | $\beta_1$ | -0.18 | 0.41 | 0.18 | 0.63 | -1.95 | 0.051 |
| | $\beta_2$ | 0.30 | 0.68 | -0.59 | 1.64 | 2.28 | 0.023 |
| | $\beta_3$ | -0.76 | 2.10 | 0.56 | 0.45 | -1.87 | 0.062 |
| | $\zeta$ | 0.00 | 0.05 | 0.01 | 0.07 | -0.62 | 0.534 |
| Perceptual model<br>trajectory condition<br>mean | $\delta_1$ | -0.01 | 0.05 | -0.01 | 0.05 | 0.08 | 0.934 |
| | $\delta_2$ | -0.02 | 0.08 | 0.04 | 0.09 | -1.58 | 0.115 |
| | $\delta_3$ | -0.01 | 0.07 | -0.01 | 0.04 | 0.50 | 0.619 |
| | $\varepsilon_1$ | -0.03 | 0.17 | -0.09 | 0.12 | 1.74 | 0.081 |
| | $\varepsilon_2$ | 0.00 | 0.09 | -0.04 | 0.04 | 2.70 | **0.007** |
| | $\varepsilon_3$ | -0.07 | 0.06 | -0.01 | 0.34 | -1.95 | 0.051 |
| | $\psi_1$ | -0.10 | 0.56 | -0.25 | 0.46 | 1.49 | 0.135 |
| | $\psi_2$ | 0.20 | 0.47 | -0.18 | 0.23 | 3.11 | **0.002** |
| | $\psi_3$ | -3.91 | 1.23 | 0.89 | 6.93 | -3.82 | **<0.001** |
| | $\sigma_1$ | 0.02 | 0.06 | -0.03 | 0.03 | 2.78 | **0.005** |
| | $\sigma_2$ | 0.20 | 0.47 | -0.18 | 0.23 | 3.11 | **0.002** |
| | $\sigma_3$ | -0.73 | 0.23 | 0.35 | 1.36 | -4.02 | **<0.001** |
| | $\mu_1$ | -0.02 | 0.12 | 0.05 | 0.13 | -1.20 | 0.229 |
| | $\mu_2$ | 0.01 | 0.42 | 0.14 | 0.38 | -0.25 | 0.803 |
| | $\mu_3$ | 0.00 | 0.15 | 0.09 | 0.18 | -1.58 | 0.115 |

IQR: Inter-quartile range
CRPS: Complex Regional Pain Syndrome
HC: Healthy control
Z: Mann-Whitney U test effect size
p: 2-tailed probability of Type I error
In bold: significant p values considering an FDR threshold for all tests in table of $p < 0.005$

### Model selection: Random effects analysis on log-model evidence

Bayesian model selection is a principled method to trade off each model’s ability to predict the observed data as a function of their complexity (which grows with the number of estimable parameters). Each model’s log-evidence was estimated from the HGF optimisation algorithm (specifically, it is the variational Bayesian approximation to the model’s marginal likelihood). Bayesian model selection was performed at the group-level using random-effects analysis (Rigoux et al., 2014), and in a family-wise manner (considering families of all models containing each perceptual and response model), using the VBA toolbox (Daunizeau et al., 2014). In a random-effect analysis, models are treated as random effects that could differ between participants, with an unknown population distribution (described in terms of model frequency with which any model prevails in the population). In addition to estimating model frequencies in the population, the VBA toolbox calculates two further summary statistics. The exceedance probability (EP, (Rigoux et al., 2014)) measures how likely it is that any given model is more frequent than all other models in the comparison set. Protected exceedance probabilities (PEPs) further correct EPs for the possibility that observed differences in model evidences (over participants) are due to chance (Rigoux et al., 2014). We used the PEP to make decisions about which model is the “best” by only selecting models with greater than 95% PEP, which corresponds to 95% confidence in that model.

In particular, we concentrated on the PEP over families of (perceptual or response) models. This involved partitioning the model space into subsets (model families), allowing reduction of the model space from 18, to two comparisons of 3 (perceptual) and 6 (response), which has advantages in preventing the over-fitting that can occur by specifying too large a model space (Rigoux et al., 2014). Each family of models contained all models that shared a common feature: either sharing the same perceptual model or same response model.

### HGF model evidence using factorial model comparison

Factorial model selection was used to select the model with the greatest log-model evidence among the four perceptual models (and summing evidence across all six response models). The greatest evidence (Supplementary Fig. 1b) was attributed to perceptual model 1, describing a single sensory noise parameter for both arms and digit pairs. Using the same factorial model selection strategy but summing over all perceptual models, the response model with the greatest evidence was number 5, which predicted trial-wise RTs from uncertainty parameters from the three levels of the HGF. The combination of perceptual model 1 and response model 5 reached exceedance probability of 90%, however, using the protected exceedance probability (PEP) there were no models that were more likely to be frequent, as the Bayesian Omnibus Risk statistic (the posterior probability that model frequencies are all equal) was a value of 1. In other words, apparent differences in model frequencies based on EP may have been due to chance. There is therefore no basis for discriminating these models based on the log-model evidence.

**Supplementary Figure 1:**
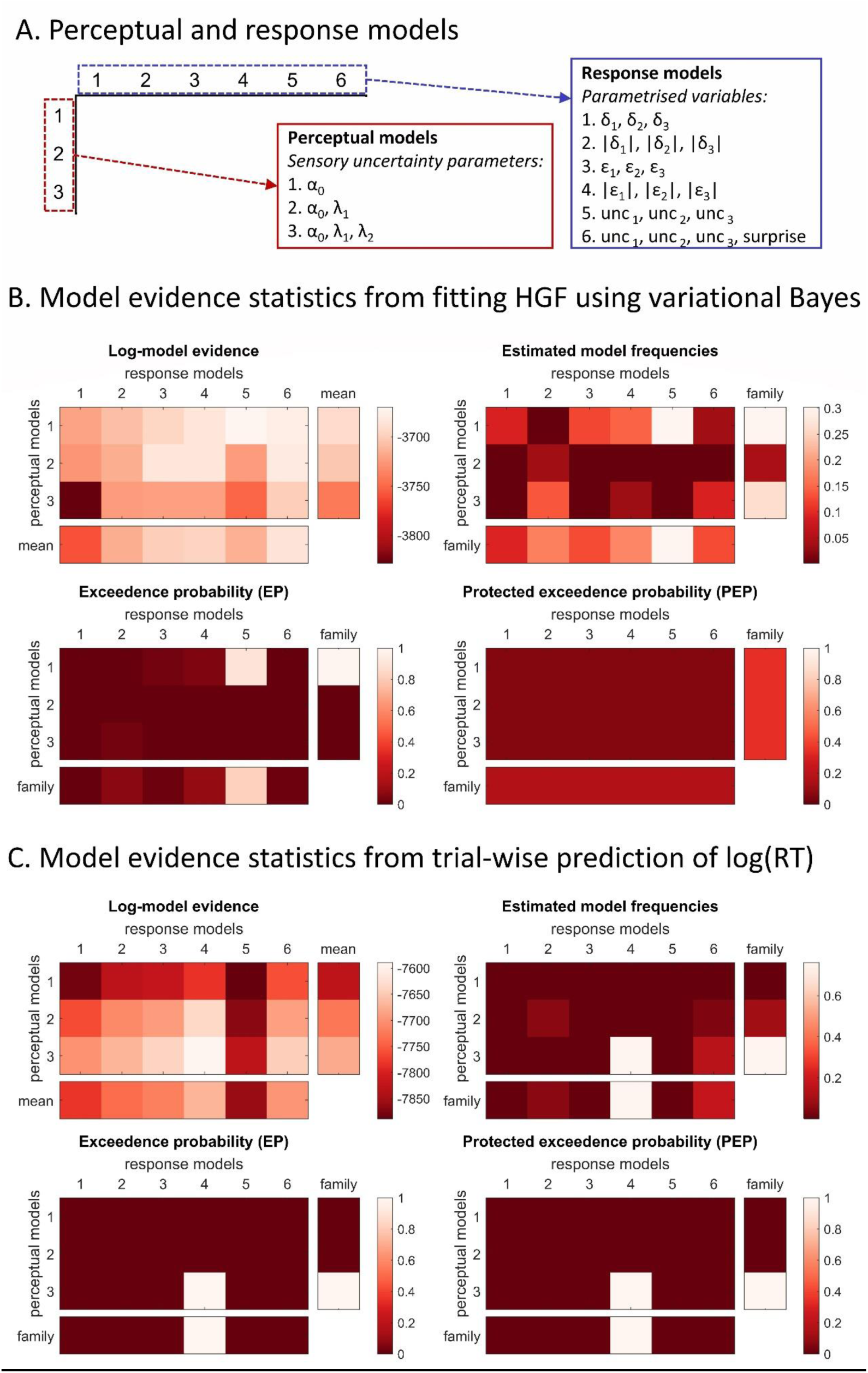
Model evidences from the two approaches used to select the best HFG model. A. Factorial model space of 3 perceptual models x 6 response models. Briefly, perceptual models differed according to whether CD (λ_1_ parameter, p-models 2 and 3) and Side (λ_2_ parameter, p-model 3) effects were considered to modify sensory uncertainty (α_0_). Response models varied according to the variables in the HGF used to predict response time, namely prediction error in its signed or absolute forms (δ and |δ|), precision-weighted precision error in its signed and absolute forms (ε and |ε|), various measures of uncertainty at each level (unc) and a measure of Bayesian surprise. B. Approximate log-model evidences (LME) from variational Bayesian estimation of each of the 18 candidate HGF models. To summarise, based on the PEP (rightmost), the best model could have arisen by chance. From the left: LME values with means shown in the right column and bottom row for each model family; Estimated model frequencies, i.e. the proportion of the time the model is expected to be represented in the population relative to the other models (values from all models sum to 1); Exceedance probability (EP), i.e. how likely it is that any given model is more frequent than all other models; Protected exceedance probability (PEP), which correct EPs for the possibility that observed differences in model evidences (over participants) are due to chance. For model frequencies, EP and PEP, the right-most column and bottom row show the estimated values using family-level inference. C. Approximate model evidences (-WAIC) from Bayesian linear regression of actual log(RT) on simulated log(RT) from the 18 candidate HGF models. See (B) legend for a description of the statistics. Using the PEP statistic, this identifies one model (perceptual model 3, response model 4) of being above-chance in its accuracy of predictions about behaviour.

### Model selection: Simulation of behaviour

Recently there has been better recognition of the importance of falsifying candidate models using simulation, to test whether the simulated outputs make accurate predictions about behaviour (Palminteri et al., 2017). Although approximations of the model evidence are useful in taking model parsimony into account, it is possible for the procedure to identify overly simplistic models that can be falsified by showing that it is unable to account for a specific behavioural effect of interest. Such models can then be rejected regardless of their log-model evidence. This “generative performance” (Palminteri et al., 2017) of the model can be assessed by simulating the model and comparing the simulated data to the observed data.

All 18 models were therefore simulated, for every subject individually, using the subject-specific parameters estimated from each model. For robustness of the findings, we use a split-half cross-validation method to test generative performance. Trial were randomly split into training (50%) and testing (50%) sets, each containing balanced numbers of trials between conditions. Parameters were estimated from the training trials, while generative performance was tested on the remaining test trials. Model predictions were then assessed in a family-wise manner for whether they were complex enough to adequately predict behaviour using the following methods:

a. The more principled method involved estimating the model evidence from a linear regression (see Fig. 2a of main paper) between the simulated log-RTs and the observed log-RTs over all trials. For each subject and model, Bayesian linear regression (Bayesreg toolbox, (Makalic and Schmidt, 2016)) was conducted with a g prior (typically used for model comparison) and a Gaussian error distribution, uses MCMC sampling of the posterior (1000 samples). This produces the WAIC approximation to the model evidence that can be used for model selection. Model selection then proceeded as described in the previous section using random-effects analysis over models, and in a family-wise manner. The model best predicting behaviour was identified using the PEP estimates for the perceptual and response model families.
b. The less principled, but most intuitive, method was based on visual inspection of the simulated data using violin plots and scatterplots. Simulated within-subject condition effects on log-RTs (namely, the two effects of change probability and digit-change distance) were visually compared to those of observed data. Furthermore, simulated individual differences in condition effects on RTs were plotted against observed differences. Spearman’s rank correlation statistics are also reported.

### Model performance results and selection via simulation

Based on visualisation of simulated behaviour (Supplementary Fig. 2a), the most striking observation was that the perceptual model (p-model 1) and response model (r-model 5) with the greater log-model evidence from variational Bayes optimisation were clearly falsified on the basis of not reproducing within-subject condition effects (Supplementary Fig. 2a) or between-subject differences in CD effects (Supplementary Fig. 2b) on RTs. In order to identify a suitable model, then, we employed Bayesian model comparison, but this time applied to the predictive accuracy of the models, i.e. how well they predict (using Bayesian linear regression) actual response times over trials (Supplementary Fig. 1c). For perceptual model 3, family-wise model comparison identified PEP of 100%. The response model with the highest predictive accuracy was the absolute precision-weighted prediction error model (model 4). For response model 4, family-wise model comparison identified PEP of 100%. These results concur with visual representations of simulated condition effects on log-RT (Supplementary Fig. 2) in which condition effects closely mirror those of actual behaviour (Fig. 3 of main paper).

**Supplementary Figure 2:**
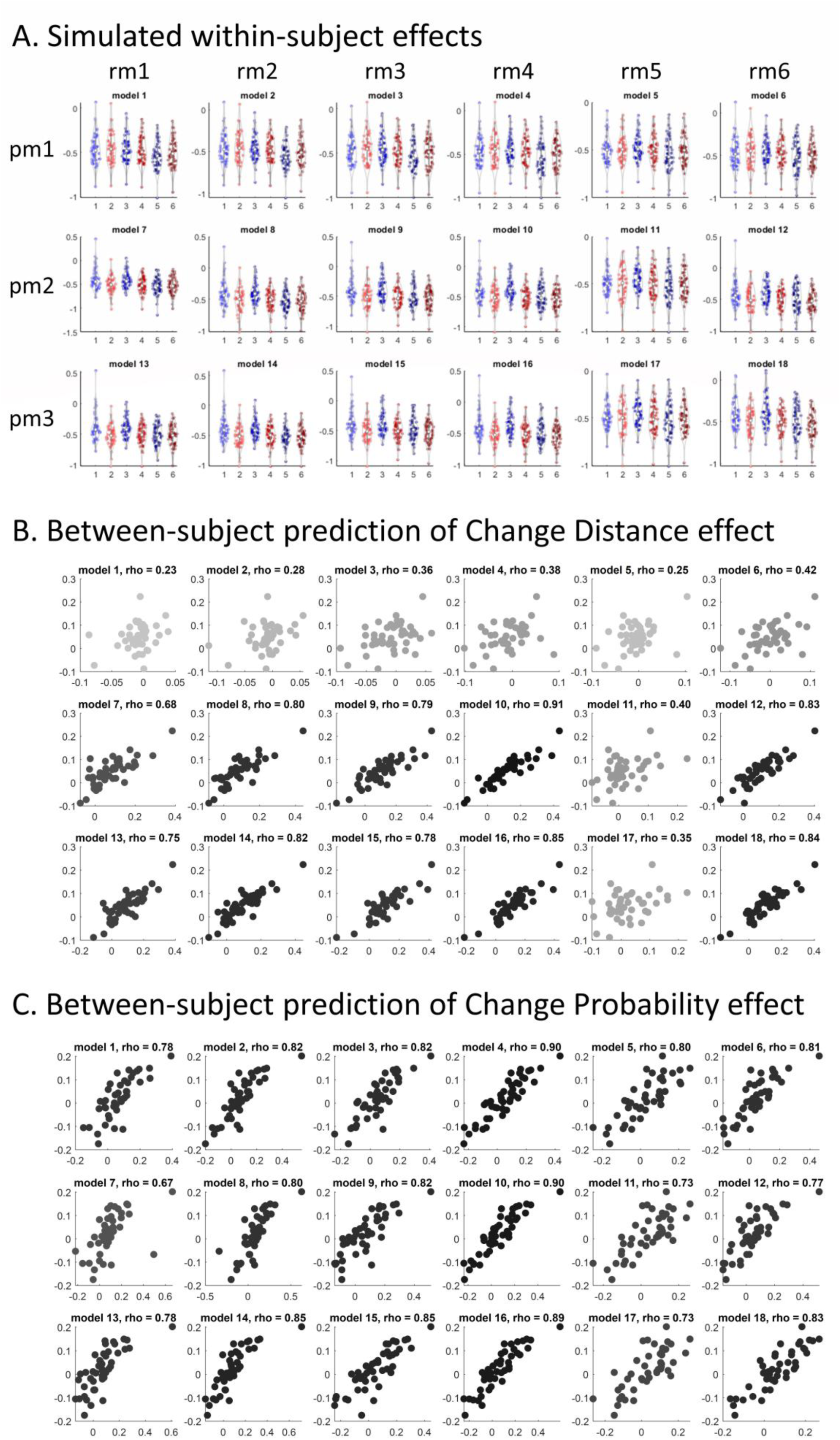
Descriptive plots of simulated log-RT data over candidate models. A. Simulated within-subject effects from each of 18 candidate models. For each model, data are simulated for each participant (individual data points on each plot) from each cell (individual violins) of the factors Change Probability (CP) and Change Distance (CD). pm: perceptual model. rm: response model B. Simulated between-subject correlation between predicted (simulated) and actual differences in log(RT) values arising from the Change Distance contrast (CD1 minus CD3, which results in increased response times). Darker shades of the data points indicated a stronger Spearman’s rho correlation coefficient. C. Simulated between-subject correlation between predicted (simulated) and actual differences in log(RT) values arising from the Change Probability contrast (10% minus 50%, which results in increased response times).

